# Getting Smaller by Denaturation: Acid-Induced Compaction of Antibodies

**DOI:** 10.1101/2022.09.19.508607

**Authors:** Hiroshi Imamura, Ayako Ooishi, Shinya Honda

**Affiliations:** Biomedical Research Institute, National Institute of Advanced Industrial Science and Technology, 1-1-1, Higashi, Tsukuba, Ibaraki 305-8566, Japan; Department of Applied Chemistry, College of Life Sciences, Ritsumeikan University, 1-1-1 Noji-Higashi, Kusatsu, Shiga 525-8577, Japan; Department of Bio-Science, Nagahama Institute of Bio-Science and Technology, 1266 Tamura, Nagahama 526-0829, Japan

## Abstract

Protein denaturation is a ubiquitous process that occurs both *in vitro* and *in vivo*. While the molecular understanding of the denatured structures of proteins is limited, it is commonly accepted that the loss of unique intramolecular contacts makes proteins larger. Herein, we report compaction of the immunoglobulin G1 (IgG1) protein upon acid denaturation. Small-angle X-ray scattering coupled with size exclusion chromatography revealed that IgG1 radii of gyration at pH 2 were ∼75% of those at a neutral pH. Scattering profiles showed a compact globular shape, supported by analytical ultracentrifugation. The acid denaturation of proteins with size reduction is energetically costly, and acid-induced compaction requires an attractive force for domain reorientation. Such intramolecular aggregation may be widespread in immunoglobulin proteins as non-canonical structures. Herein, we discuss the potential biological significance of these non-canonical structures of antibodies.

## Introduction

Proteins are polymer chains composed of amino acid residues and optimised sequences that have the distinct thermodynamic states,^[1]^ one of which is a native state and another a denatured state. The folded structure in the native state has been determined by extensive biophysical investigations and predicted via bioinformatic methods. In contrast, the molecular pictures for the structures in the denatured state are limited. The denatured state comprises a variety of dynamic conformers, including fully and partially unfolded structures, which has technically hindered the elucidation of their fine structures. Despite this, recent studies have explored the difference in the denatured structure causing different biological and immunological consequences; for example, an expanded conformation of the Tau protein is favourable for biomolecular condensation, which promotes amyloid formation.^[2]^ Whether unfolded structures of a protein (PNt) are expanded or collapsed determines the efficiency of the secretion to the cell surface.^[3]^ Reversal effects of therapeutic proteins are potentially associated with the morphology of the denatured aggregates.^[4,5]^ These studies indicate the significance of accumulating knowledge about the denatured structures of proteins.

Both amino acid side chains and main chains stabilise the native protein structure by forming non-covalent bonds among them, where the constituent atoms are uniquely organised. Upon denaturation, these native contacts are replaced by non-specific water-protein contacts that increase the spatial and temporal dynamics of the proteins.^[6]^ Thus, the size of proteins is always increased by denaturation,^[7-9]^ which is a major principle of protein folding.

Here, we report an exceptional case where denaturation makes proteins smaller via an immunoglobulin G1 (IgG1) monoclonal antibody. Acid-solution induces the denaturation of IgG1, including the loss of the binding function^[10]^ and native structures.^[11,12]^ However, the structures of the denatured IgG1 remain unclarified. This study used a small-angle X-ray scattering (SAXS) technique^[13]^ to demonstrate the size reduction of IgG1 by probing the global solution structure of the antibody under neutral and acidic pH conditions. While the biological significance of the reduced size of denatured IgG1 was unclear, the current models include the molecular requirements of non-canonical contacts between domains, suggesting the universality of the phenomenon of denaturation reducing the size of immunoglobulin proteins.

## Results

Figure 1 depicts the acid-induced changes in an IgG1 monoclonal antibody, mAb-A, when measured by SAXS. The SAXS profiles were similar at pH 7.1–3.1, with the characteristic shoulders in the log–log plots (Figure 1a, arrows) and peaks in the Kratky plots (Figure 1b) at *q* = ∼0.08 and ∼0.16 Å^−1^. The former trait (*q* = ∼0.08 Å^−1^) corresponds to the Fc (crystallizable fragment)–Fab (antigen-binding fragment) or Fab–Fab distance (∼80 Å). The latter trait (∼0.16 Å^− 1^) is sensitive to the inter-domain distances/orientations between the adjacent domains.^[14]^ These scattering patterns share these characteristics with the theoretical scattering calculated from the crystal structure of IgG1 (black line in Figure 1; denoted as “Model”) and previous SAXS data reported in the literature.^[15,16]^ Conversely, the scatterings at pH < 2.0 were distinct from those at pH 7.1–3.1; the characteristic shoulders (Figure 1a) and peaks (Figure 1b) diminished at pH < 2.0, with more scattering in the low-*q* range at pH 1.1 (Figure S1a).

**Figure 1.**
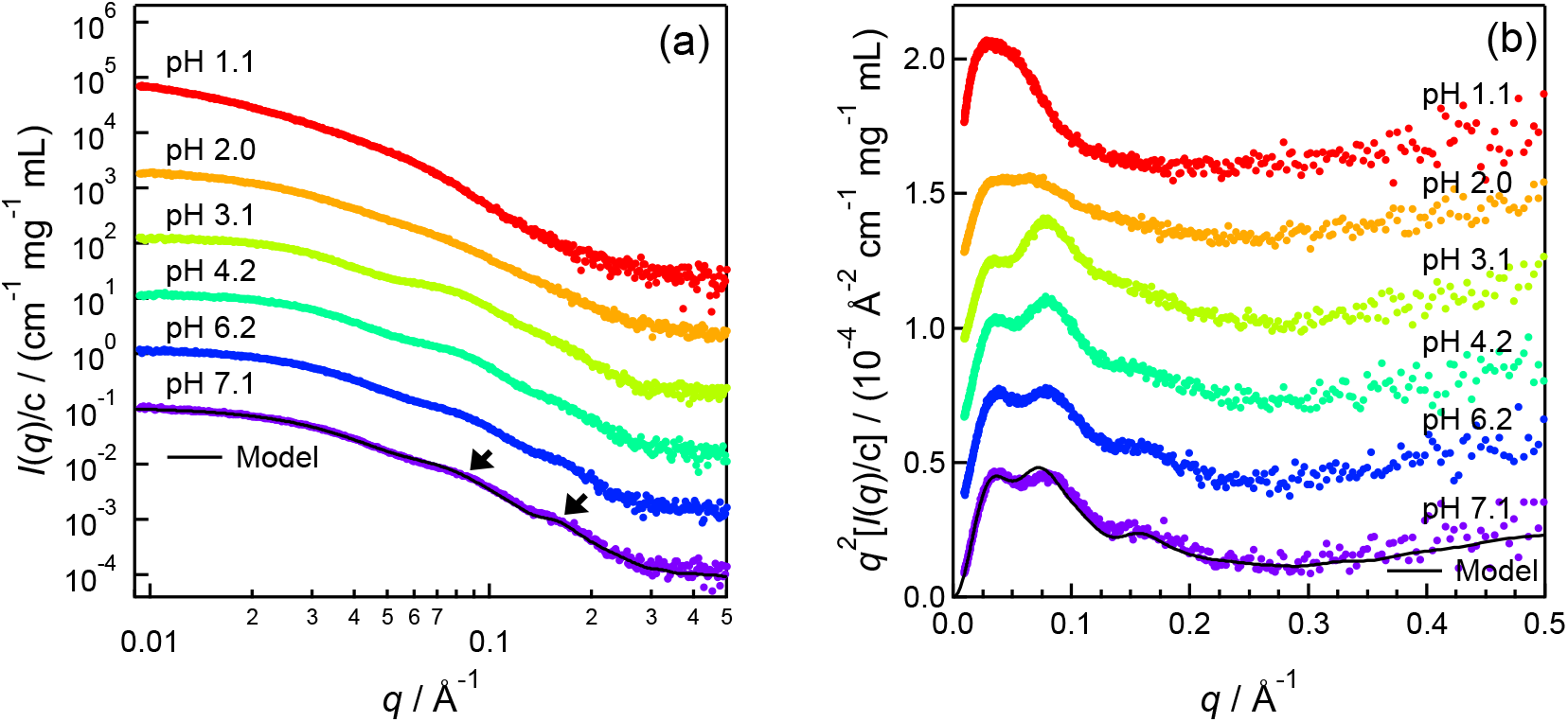
The mAb-A pH-dependence of the small-angle X-ray scattering (SAXS) at 25 °C. Concentration-normalised SAXS profiles are presented as (a) log–log and (b) Kratky plots. The protein concentration was 3.5 ± 0.1 mg/mL. The data are vertically shifted for clarity. The black line shows the theoretical SAXS profile of the IgG1 model (PDB: 1HZH). The arrows in (a) indicate the shoulders at *q* = ∼0.08 and ∼0.16 Å^−1^.

The SAXS data indicated a drastic conformational alteration of mAb-A around pH 2, but the contaminating aggregates hampered further structural analysis (see Supporting Information and Figure S1). Therefore, we performed SEC-SAXS analysis^[17-19]^ *in situ* SAXS during size exclusion chromatography (SEC), which allowed us to monitor the SAXS of monomeric mAb-A at pH 2. Chromatographic analysis was conducted of the integral SAXS intensities and ultraviolet (UV) absorbance at 280 nm using an elution buffer solution of 0.1 M glycine-HCl (pH 2) with 0 M or 0.2 M NaCl (Figure 2a). The peaks at ∼169 min (0 M NaCl) and ∼66 min (0.2 M NaCl) represented monomeric mAb-A that was well separated from the aggregates, while the peaks at ∼124 min (0 M NaCl) and ∼56 min (0.2 M NaCl) indicated dimeric mAb-A. The aggregates (> trimer) manifested as peaks that presented early in the analysis. These assignments are supported by the chromatogram of the integrated SAXS intensities, which have more pronounced aggregates than the UV absorbance. The variation in the retention time between the data (0 M and 0.2 M NaCl) was due to the difference in the flow rates used.

**Figure 2.**
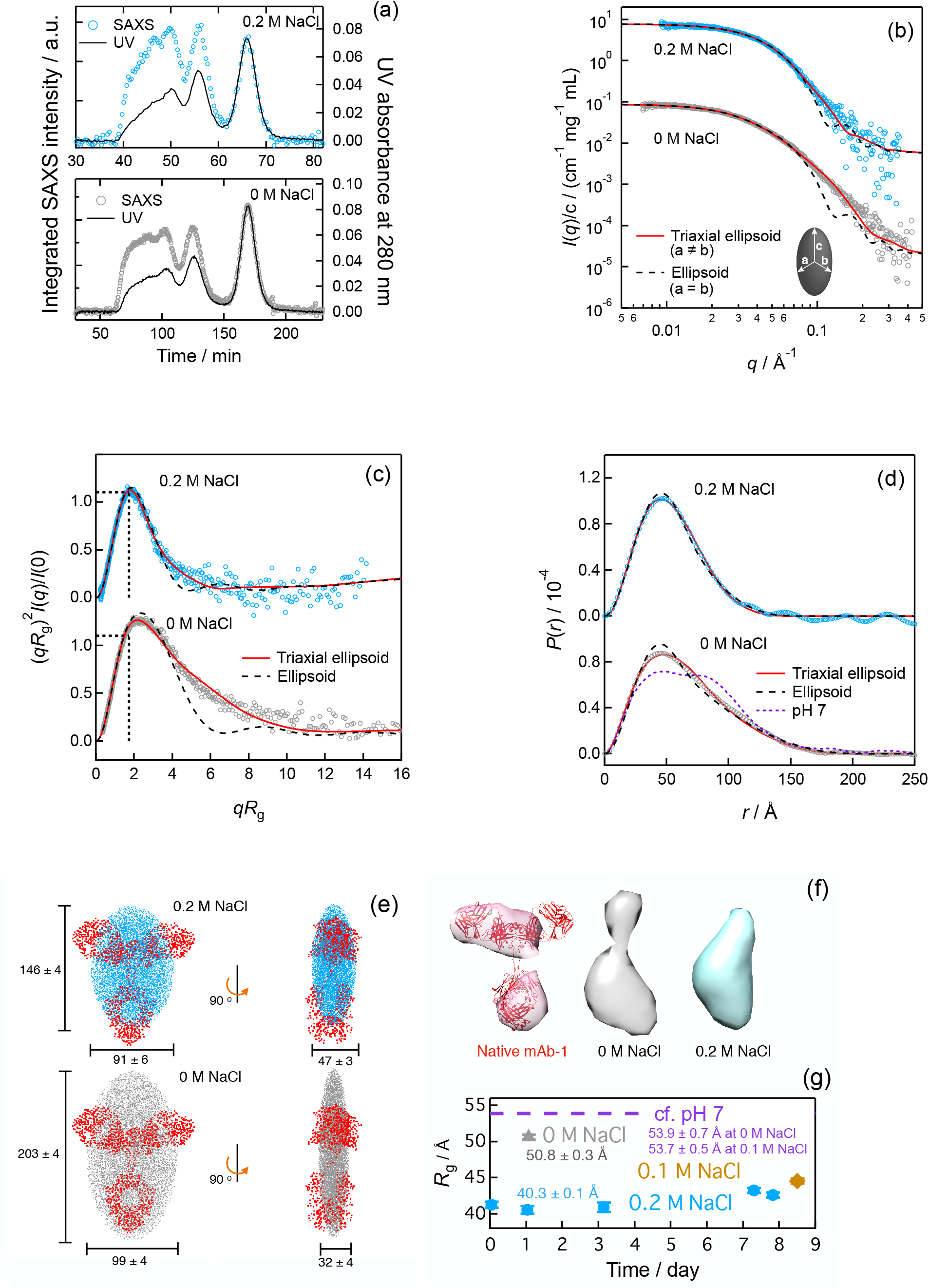
Size exclusion chromatography small-angle X-ray scattering (SEC-SAXS) analysis on mAb-A at pH 2. The results of mAb-A at 0 M NaCl and 0.2 M NaCl are represented as grey and cyan, respectively. (a) Overlaid elution SAXS and UV profiles where the SAXS-derived chromatogram is an integrated SAXS intensity vs. retention time. The elution UV profiles were acquired by simultaneous recording of mAb-A absorbance at 280 nm. (b) SAXS profiles of monomeric mAb-A. The scattering for mAb-A in 0.2 M NaCl was multiplied by 10^2^ for clarity. Theoretical scattering of the triaxial ellipsoid (red solid line) and ellipsoid (dashed black line) with best fit parameters for mAb-A are shown. An elliptical object indicates where *a, b*, and *c* are the length parameters. (c) Dimensionless Kratky plots of IgG1 SAXS profiles and theoretical scattering. (d) The distance-distribution function, *P*(*r*), by direct Fourier transformation of the SAXS profiles. *P*(*r*) of the triaxial ellipsoid (red solid line) and ellipsoid (black dashed line) were calculated from the scatterings in (b). *P*(*r*) of the native mAb-A at pH 7.1 (Figure S1e) is overlaid for comparison. This *P*(*r*) was scaled such that the area is equivalent to the *P*(*r*) of the mAb-A at pH 2 and 0 M NaCl. (e) A point cloud representation of the triaxial ellipsoids using best fit parameters for the scatterings of mAb-A at 0 M NaCl and 0.2 M NaCl. A canonical IgG1 structure,^[26]^ where the C_α_ atoms (red point clouds) are placed to fit with the triaxial ellipsoids with characteristic lengths. (f) 3D reconstructed models of the mAb-A at pH 2 using *ab initio* electron density determination (DENSS).^[25]^ The canonical mAb-A models are also shown, indicating the *ab initio* model reconstructed from the SAXS data (light pink), overlaid with the atomic coordinates of the IgG1 (red ribbons).^[26]^ All the volumes inside the model surfaces are set at 258,000 Å_3_, where the Porod volume was determined for the scattering of mAb-A at 0 M NaCl. (g) Dependence of mAb-A *R*_g_ at pH 2 on incubation time and NaCl concentration. The *R*_g_ of the native mAb-A at pH 7 (with 0 M NaCl and 0.1 M NaCl) is indicated for comparison.

Based on the chromatograms and their frame-by-frame analyses (Figure S2), we determined the concentration-normalised SAXS profiles (*I*(*q*)/*c*) of monomeric mAb-A at pH 2 (Figure 2b). Because the concentrations were < 0.5 mg/mL in the sample cell during the chromatographic analysis, the intermolecular interference was negligible. The radius of gyration (*R*_g_) was 50.8 ± 0.3 Å for 0 M NaCl and 40.5 ± 0.4 Å for 0.2 M NaCl, and the concentration-normalised zero-angle scattering (*I*(0)/*c*) was 0.0894 ± 0.0003 cm^2^ mg^−1^ (122 kg/mol) for 0 M NaCl and 0.0783 ± 0.0004 cm^2^ mg^−1^ (108 kg/mol) for 0.2 M NaCl, which were comparable with the native monomeric mAb-A of 0.0880 ± 0.0007 cm^2^ mg^−1^ (121 kg/mol). The comparable molar masses (Table S1) were found via Bayesian inference^[20]^ These results indicated that the SEC-SAXS was able to capture the monomeric acid-denatured mAb-A by removing the aggregates. The deviations from the exact molar mass (*M*) of mAb-A (148 kg/mol) were due to a systematic error. In the log–log and Kratky plots of the SAXS profiles (Figure 2b and S2h), the characteristic shoulders/peaks observed for the native mAb-A at pH 7.1 (Figure 1) was not observed at pH 2. The distance-distribution function (*P*(*r*)) at pH 2 indicated a lack of the characteristic distance distribution (∼80 Å) observed at pH 7.1 (Figure 2d), suggesting the distances and orientations of the Fab-Fc and Fab–Fab regions were not canonical at pH 2. Furthermore, the maximum distance (*D*_max_) at pH 2 were < ∼200 Å (0 M NaCl) and < ∼150 Å (0.2 M NaCl). The former was comparable with *D*_max_ at pH 7.1.

A fitting analysis of the SAXS profiles characterised the shape of mAb-A at pH 2 (Figure 2b). Among the theoretical scattering functions of the sphere, cylinder, elliptical cylinder, ellipsoid, and triaxial ellipsoid shapes,^[21,22]^ the scattering function of the triaxial ellipsoid (*a* ≠ *b*) was the best approximation, with lengths, *a, b*, and *c* defining the shape of the triaxial ellipsoid (Figure 2b). The scattering function of the ellipsoid (*a* = *b*) produced deviations from the experimental scatterings in the region of *q* > 0.1 Å_–1_ in Figure 2b and *qR*_g_ > 4 in Figure 2c; such a deviation was also found in *P*(*r*) functions (Figure 2d). These deviations for *I*(*q*)/*c* and *P*(*r*) at 0 M NaCl were larger than those at 0.2 M NaCl, indicating that the shape of mAb-A at 0 M NaCl was more flattened than that at 0.2 M NaCl.

Figure 2c shows the dimensionless Kratky plots. The SAXS of an ideal globular protein obeys Gunier’s law, *I*(*q*) = *I*(0)exp(–(*R*_g_*q*)^2^/3), and shows the maximum value where the black dotted lines intersect in the dimensionless Kratky plots:^[23,24]^ *qR*_g_ = 3^1/2^ and (*qR*_g_)^2^*I*(*q*)/*I*(0) = 3exp(– 1) = 1.104. The maximum peak, 0.2 M NaCl, of the dimensionless Kratky plot indicates that the shape of mAb-A closely resembles that of the ideal globular proteins. Slight deviation of the maximum at 0 M NaCl from the “ideal” maximum would be due to the flattened ellipsoidal shape, although the relevance thereof to conformational flexibility could not be rejected.

Figure 2e visualises the triaxial ellipsoidal approximation with the best fit parameters for the scatterings of mAb-A. The representation of mAb-A at pH 2 with 0 M NaCl (grey) and 0.2 M NaCl (blue) was flattened, in comparison with that of the native IgG1 (red). We observed that 0.2 M NaCl made mAb-A more compact with a smaller maximum length and globular shape than 0 M NaCl. Figure 2f shows the reconstructed 3D-models of mAb-A at pH 2 from the SAXS profiles and *P*(*r*) by the *ab initio* electron density determination, DENSS (DENsity from Solution Scattering), algorithm.^[25]^ The flattened shapes of mAb-A at pH 2 were also consistent with those obtained via the fitting analysis that used the theoretical scattering functions. In addition, these 3D models illustrated some anisotropic electron distributions, such as a lower density region that looked like a “neck” in 0 M NaCl. Another remarkable characteristic of mAb-A at pH 2 included a smaller *R*_g_ than that of the native mAb-A (Figure 2g). The addition of NaCl led to a smaller *R*_g_ at pH 2, while *R*_g_ was not dependent on NaCl at pH 7. mAb-A at pH 2 was very plastic and presented as globular due to the shielding effect of the salt, which reduced the intramolecular electrostatic repulsion of the acid-denatured mAb-A. We noted that the incubation time at pH 2 did not significantly change the *R*_g_.

We further conducted SEC-SAXS analysis on another IgG1, mAb-B, to test whether a variation of the sequence in the variable region alter the conclusion of the size reduction. We verified an acid-induced compaction of mAb-B, which showed *R*_g_ = 53.0 ± 0.4 Å at pH 7.2 and *R*_g_ = 46.5 ± 0.3 Å at pH 2 and 0.2 M NaCl.

An orthogonal analysis of the acid-denatured mAb-A using an analytical ultracentrifugation (AUC) strengthened the SAXS analysis. The sedimentation velocity data (Figure S3a) showed the distribution of the sedimentation coefficients (*s*) of the mAb-A samples at pH 7 and pH 2 (Figure S3b), in which prominent peaks at *s* = ∼6.5–7.0 S were consistent with the expected *s* value for the known molar mass of monomeric mAb-A (148 kg/mol; Table S2). We found a small but significant variation in *s*_20,w_ (*s* at 20 °C in water) among the samples, where the effects of the viscosities and densities of the solvents were corrected. This variation indicates the differences in the molecular envelope, represented as the friction factor in the AUC. The variation in *s*_20,w_ due to the association of the molecules was ruled out; for example, the dimer of mAb-A had a *s* = ∼9.1 S at pH 2 and 0 M NaCl (Figure S3b). According to equations (S8–S11), the friction factor (*f*/*f*_0_) and the corrected friction factor (*f*/*f*_0,sp_) were determined. The *f*/*f*_0_ of the native mAb-A (1.43) was comparable with a previously reported value (1.55);^[27]^ the deviation should be due to the difference in the specific volume (*v*) used. Empirically, approximately globular proteins, moderately elongated proteins, and highly elongated proteins were reported to display *f*/*f*_0_ of 1.2–1.3, 1.5–1.9, and 2.0–3.0, respectively.^[28]^ The current *f*/*f*_0_ indicated that the native mAb-A and the acid-denatured mAb-A were both globular rather than elongated. The *f*/*f*_0_ at pH 2 decreased from 1.58 to 1.35 with the addition of 0.2 M NaCl, consistent with the SAXS result that mAb-A at pH 2 was more globular in 0.2 M NaCl. The corrected friction factor (*f*/*f*_0,sp_) in equations (S9–S11) provided information regarding the protein shape and can be calculated for any triaxial ellipsoid.^[29,30]^ We calculated *f*/*f*_0,sp_ using the triaxial ellipsoid lengths *a, b*, and *c*, as determined using the SAXS analysis (Figure 2). The experimentally determined *f*/*f*_0,sp_ at pH 2 decreased by 0.19 (1.32 at 0 M NaCl and 1.13 at 0.2 M NaCl), which was comparable with the decrease in the calculated *f*/*f*_0,sp_ by 0.14 (Table S2). The differences between the absolute values of the experimental *f*/*f*_0,sp_ and the calculated value were due to uncertainties in the *β*_1_, *v*, and *η* used in equations (S7 and S8). Together, the results of SAXS—the non-elongated shape of the acid-denatured mAb-A and its plastic globularisation by the addition of NaCl— are well supported by the AUC.

## Discussion

Protein denaturation increases the size of proteins independent of perturbations such as temperature, pressure, or chemical denaturation.^[8]^ Indeed, the *R*_g_ for proteins reported in the literature indicated that the underlying power laws of *R*_g_ vs. chain lengths (*L*) are distinct between native and unfolded protein structures (Figure 3). The *R*_g_ of the native state of the multidomain protein mAb-A followed the power–law relationship for the native structures, although the representation of *R*_g_ in the literature^[9]^ is dominated by single-domain proteins. In contrast, the *R*_g_ of non-native mAb-A was not predicted by the power–law relationship and anomaly smaller than that of the native state. The *R*_g_ of the non-native mAb-A dimer, apart from the scaling law for unfolded proteins, also indicates a non-random coil structure.

**Figure 3.**
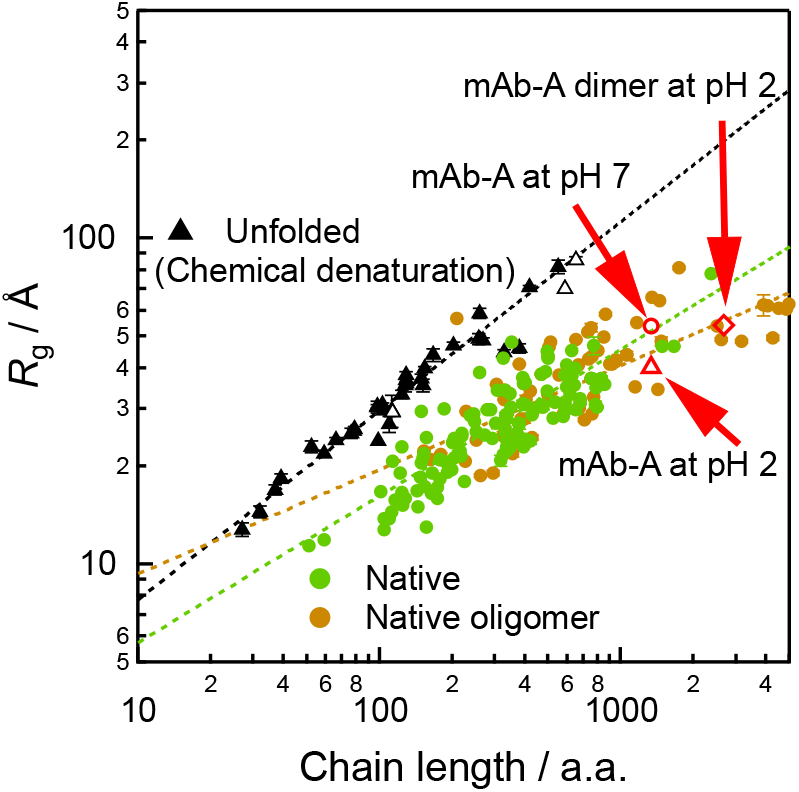
*R*_g_ dependence on protein chain length. The data in literature includes native (green, closed circles), native oligomeric (dark orange, closed circles), and unfolded (black triangles, where closed and open triangles indicate disulfide-free and disulfide-bonded) proteins, curated by Po-Min Shih *et al*.^[9]^ The dotted lines represent the power–law scaling of *R*_g_ = *R*_0_*L*^*ξ*^, where *R*_0_ is a prefactor (2.04, native; 4.47, native oligomer; and 2.05, unfolded), *L* is the amino acid (a.a.) chain length, and ξ is the scaling exponent (0.45, native; 0.32, native oligomer; and 0.58, unfolded). *R*_g_ of mAb-A at pH 7 (red, open circle), *R*_g_ of mAb-A at pH 2 with 0.2 M NaCl (red, open triangle), and *R*_g_ of mAb-A dimer at pH 2 with 0.2 M NaCl (red, open diamond) determined in this study are overlaid onto previously reported results.^[9]^ The *R*_g_ value of mAb-A at pH 7 is given as an infinite dilution of the protein (Figure S1h).

Protein acid denaturation is caused by the repulsion of positive charges on the surface of the protein that are approximated by simple models such as the Linderstrøm–Lang smeared charge model.^[31,32]^ This model indicates that the size expansion decreases the Gibbs energy by denaturation (see Supporting Information) and describes how acid denaturation that reduces protein size is energetically costly and anomalous. No previous reports have shown acid denaturation without size expansion; even the molten-globule state of cytochrome *c* at pH 2 and

0.1 M NaCl (17.4 Å), which is described as a “compact” denatured state, is larger than that at its native state (13.5 Å).^[33]^

The charge distributions shown in Figure 4a identified a positive potential on the surface of mAb-A that widely emerged at pH 2. We hypothesise that alternative, energetically favourable interactions hold the parts of the acid-denatured IgG1 molecule together by compensating for the energetic cost of maintaining the size of the protein. Additional van der Waals interactions would assist in this process. The acid-denatured state at pH 2 was more compact with the addition of NaCl, indicating that shielding the coulomb repulsion among the charged residues and unfavourable salt-induced solvation of the exposed hydrophobic protein surface assisted the sub-complete packing of the protein. These effects could be common for inter- and intra-molecular interactions. Lower pH increases intermolecular repulsive interactions, which is depicted as the intermolecular interference (Figure S1f-i). Counteracting favourable interactions between the monomers could generate the aggregates at pH 2.

**Figure 4.**
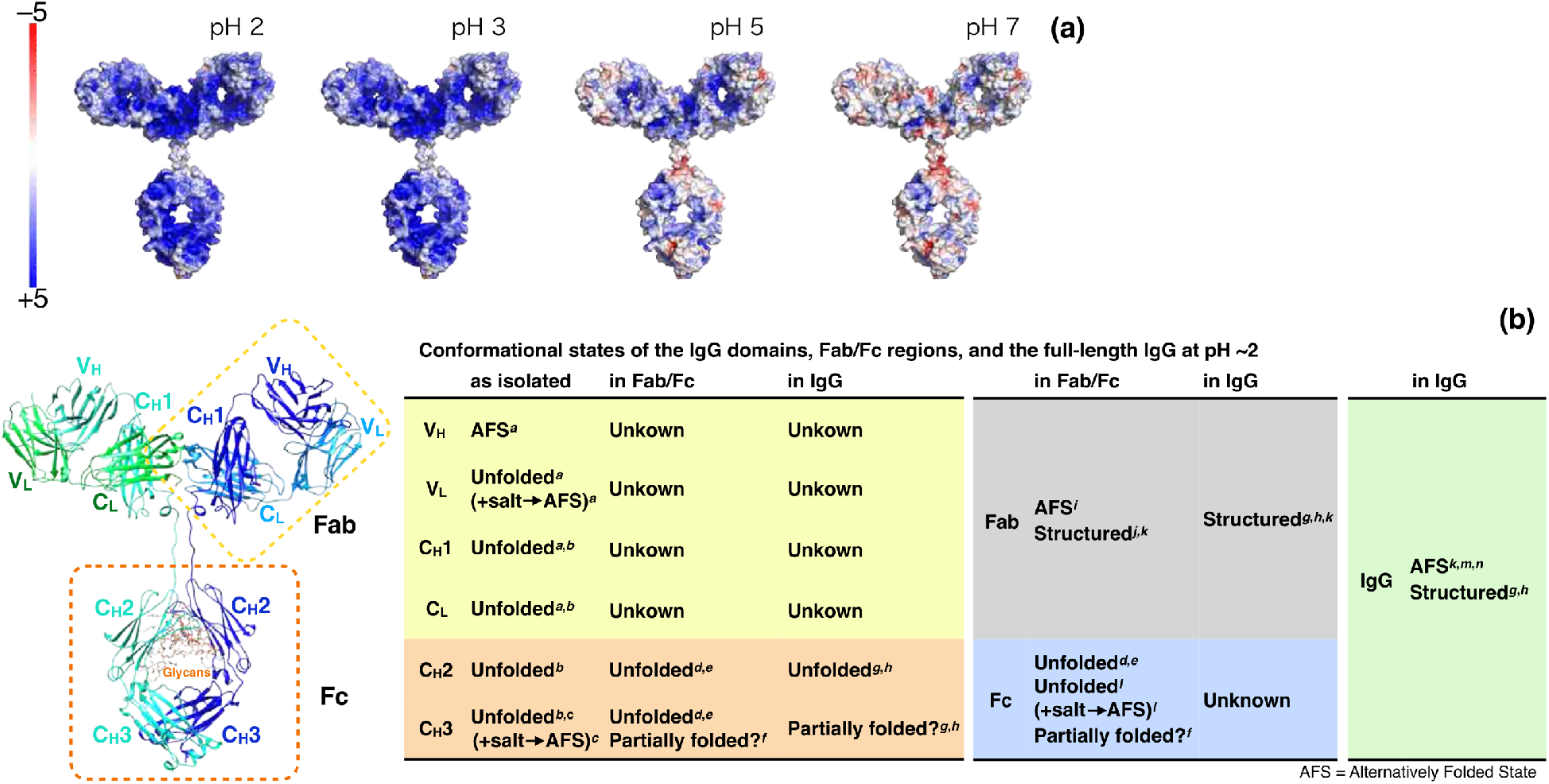
Acid effects on IgG structure. (a) Charge distribution in IgG1 at various pHs. The electrostatic properties were calculated using the p*K*_a_ predictor PROPKA^[34,35]^ and the adaptive Poisson–Boltzmann solver (APBS) software^[36]^ using the atomic coordinates for IgG1^[26]^ and the mAb-A sequence. (b) The conformational states of the IgG domains at pH 2.0–2.6 (with/without salts) in the context of the isolated domain, Fc/Fab regions, the full-length IgG reported in the literature; the references are presented in the table are ^*a*^pH 2.0;^[37] *b*^aglycosylated C_H_2, pH 2;^[38] *c*^pH 2;^[39] *d*^aglycosylated Fc, pH 2.6;^[40] *e*^aglycosylated Fc, pH 2.5;^[41] *f*^glycosylated Fc, pH 2.6;^[40] *g*^pH 2;^[42] *h*^pH 2.0;^[43] *i*^pH 2;^[44] *j*^pH 2.5;^[45] *k*^pH 2.1;^[46] *l*^glycosylated Fc, pH 2.0;^[47] *m*^pH 2.0;^[11]^ and ^*n*^pH 2.0.^[12]^ The canonical IgG1 structure,^[26]^ the domains thereof (C_H_3, C_H_2, C_H_1, C_L_, V_H_, and V_L_), and the regions (Fc and Fab) are presented.

The gain in native contacts in folded proteins contributes to their size reduction.^[7]^ Therefore, the acid-denatured state for mAb-A could be regarded as an alternatively folded state (AFS) that had alternative contacts that prevented expansion. Buchner *et al*.^[48]^ referred to AFS hitherto as “The antibody conformation at low pH had been termed ‘alternatively folded state’ because it exhibits characteristics of the folded state (e.g. remarkable stability against unfolding), but the available spectroscopic information suggests that it is structurally significantly different from the native state. It was first described for a complete IgG1 antibody,^[11]^ but single domains such as C_H_3 also can adopt this state.^[39]^” The original paper published in 1991^[11]^ characterised MAK33, a murine *κ*/IgGl subtype, at pH 2. The *s*_20,w_ of 5.53 for the AFS (pH 2) of MAK33 was smaller than the *s*_20,w_ of 6.60 for the native protein, and much larger than *s*_20,w_ of 3.2 for the denatured protein in 6 M guanidine hydrochloride at a pH 2. These AUC results indicated that the molecules were neither significantly aggregated nor fully extended. Nevertheless, the small amount of the aggregates functioned as impurities hindered the discovery that the AFS state of IgG1 can be smaller than the native state. As the SEC-SAXS and the sedimentation coefficient distribution analysis for the AUC have not been established previously, the technical limitations also hindered the discovery. The circular dichroism (CD) spectra of the AFS state of MAK33 at pH 2 were similar to the current CD spectra (Figure S4a), further supporting the AFS of the present acid-denatured mAb-A in which the β-sheets are dominant (Table S3) and slightly reorganised, as indicated using infrared (IR) analysis (Figure S4c). Subsequent studies on AFS focused on the isolated protein domains such as C_H_3 or V_L_ domains, rather than full-length antibodies. Buchner *et al*.^[49]^ reported that the MAK33 V_L_ domain could form stable high-molecular weight oligomers at pH 2 that had previously been reported as AFS. Although the term AFS is loosely defined, it can be concluded that the association between protein domains is a common mechanism for AFS irrespective of whether full-length antibodies or isolated domains are evaluated.

Studies have also reported that other IgG molecules (murine monoclonal IgG2a antibody CB4-1^[46]^ and monoclonal chimeric antibody IgG1 BR96^[12]^) can adopt the AFS state at pH 2. In addition, Sedlák *et al*^[50]^ proposed an inter-region or inter-domain interaction of heat-denatured IgG, referred to as “intramolecular aggregation”. The authors of that study compared differential scanning calorimetry (DSC) thermograms of full-length IgG and its Fab portion. The difference in the melting temperatures of the Fab domain suggested a Fab–Fab interaction in the heat-denatured state of the protein, which was supported by the lack of protein concentration-dependence in the thermograms.^[50]^ The concept of “intramolecular aggregation” would be universal for IgG irrespective of the denaturing perturbations (e.g., acid or heating).

We also investigated the domains that participated in inter-domain interactions in acid-denatured IgG1. The canonical IgG1 protein consists of 12 domains in the Fc or Fab regions (Figure 4b). According to the literature,^[11,12,37-46,51]^ the stabilities of the domains depend on the molecular contexts (Figure 4b), e.g., isolated domains or domains in a full-length IgG. Isolated C_H_3^[38]^ at pH 2 and C_H_3 in an aglycosylated Fc^[41]^ at pH 2.5 unfold; hence, the native dimerised C_H_3–C_H_3 interface is lost.^[38,41]^ However, the ordered structure of C_H_3 in the full-length IgG is possibly retained^[42,43]^ with local disorder that decreases the left-twisting of β-strands (Figure S4 and Table S3). The C_H_2 unfolds at pH 2 regardless of the presence of glycans in any molecular context.^[38,40-43]^ The isolated domains (C_H_1, C_L_, and V_L_) of the Fab region unfold at pH 2,^[37]^ whereas the Fab region is structured when the Fab region protein^[44-46]^ and full-length IgG^[42,43,46]^ protein are examined. Therefore, the possible mechanism of acid-induced intramolecular aggregation of IgG1 is as follows: at pH 2, C_H_3 avoids complete unfolding by interacting with the Fab region, where the flexibility of the unfolded C_H_2 does not hamper the inter-domain contacts. Salt weakens the Fab–Fab repulsion, which brings the two Fab regions close to each other and makes IgG1 smaller. Self-interactions between C_H_3 are stabilised by salt, indicated by the AFS of the isolated C_H_3 oligomer.^[39]^ We speculated that the hydrophobic V_H_–V_L_ and C_H_1–C_L_ interfaces^[52]^ would be exposed upon breaking the native interfacial contacts and the inter-domain salt bridges.^[53]^ All those effects could make non-canonical contacts between the Fab and Fc regions.

A lack of atomistic modelling or the ensemble representation thereof is a limitation of this study. A flexible hinge in IgG allows for it to enable diverse positioning of the Fab and Fc regions. Tian *et al*. ^[16]^ used this ensemble of models to improve the fitting of a native IgG1 SAXS profile instead of using a single model. The possible distribution of *R*_g_ of the native IgG1 was ca. 45–58 Å. This indicates that the minimal requirement for getting smaller is the non-canonical positioning of the Fab and Fc regions, which never populate in the native state. The acid-induced structural changes, manifested as the decline of the Kratky peak at *q* = 0.16 Å^−1^ (quaternary structure), and the CD/IR spectral change (secondary structure) make the irregular Fab/Fc arrangements energetically favourable. The present shape approximation of the acid-denatured mAb-A using the theoretical scatterings of the defined shapes and the DENSS method does not rule out the ensemble of conformations that is compatible with the SAXS profile but represents the dominant structures. It is intriguing to determine whether this non-canonical structure of the acid-denatured IgG1 functions similarly to an α-lactalbumin in the molten globule state (human α-lactalbumin, HAMLET), which is lethal to tumour cells.^[54]^ A few reports have shown the biological significance of the AFS of immunoglobulins; for example, the oral administration of Ig (IgG and IgA) has been reported as a passive immunity treatment. Orally administered Ig is exposed to the acidic pH of the gastrointestinal (GI) tract but is resistant to complete digestion.^[55]^ Jasion and Burnett^[11]^ suggested that transient exposure of IgG to the acidic conditions in the stomach may increase the stability of IgG as it passed into the GI tract and against proteolytic enzymes.^[55]^ The non-canonical structure of acid-denatured IgG1 presented herein supports the idea that reducing the flexibility of locally unfolded regions via intramolecular aggregation is beneficial for the protein’s resistance to enzymatic digestion. The ions present in physiological conditions could also support the tight packing of the acid-denatured IgG1 (Figure 2). Another example regards the expression and secretion of natural or recombinant Ig proteins: Onitsuka *et al*.^[56]^ found that antibody-producing Chinese hamster ovary cells secreted aggregates of recombinant IgG from the intracellular environment into the extracellular milieu, further evidenced by a new live–cell imaging technique^[57]^ that uses a fluorescently labelled AF.2A1 as a protein probe for non-native IgG structures.^[51,58]^ Similar intracellular particles, known as Russell bodies, are found in plasma cells. Russell bodies are natural inclusion bodies usually found in cells that overexpress Ig. Mammalian cells generally secrete only correctly folded proteins; therefore, a naturally occurring AFS could pass through an intracellular quality control system, such as endoplasmic reticulum (ER)-associated degradation.^[56,57]^ If the naturally occurring AFS was similar to the acid-denatured IgG1, the compact form may be favourable for Ig intracellular transport, secretion, internalisation, and transcytosis in various circumstances.

The acid-denatured state of IgG1 is relevant to downstream processes of biopharmaceutical manufacturing where antibodies experience temporal exposure to acid during affinity chromatographic purification and virus inactivation, which is followed by neutralisation. The non-native conformation of IgG1 induced by this pH shift is aggregation-prone at a neutral pH, with structural origins that remain unclear.^[10,59-64]^ Because the acid-denatured state is unstable at a neutral pH, the IgG1 molecules transition toward native or aggregated states. It is unclear whether the molecular characteristics of the acid-denatured IgG1, such as intramolecular aggregation, slow or accelerate these reactions. In addition, the precise prediction of antibody stability in an acidic environment requires the characterisation of acid-denatured structures, including in the dimeric forms (Figure S5), which is important aspect of the engineering process of antibodies.^[65]^

## Conclusion

The current knowledge-based principle of proteins getting larger by denaturation that has been established by many studies using single-domain proteins should be reconsidered. New view should now include the potential for proteins to get smaller by denaturation, as we demonstrated for acid-denatured IgG1 proteins (mAb-A and mAb-B). The reported cases are limited; nevertheless, as they shared the characteristics of the previously reported AFS for IgGs, an acid-induced compaction can be common among various IgG proteins. Multidomain proteins possess additional higher-order structures, such as domain–domain spatial arrangements or orientations, which are lost upon denaturation. These degrees of freedom in multidomain proteins are prerequisites for a more compact structure by the rearrangement of the domains, in which there are minor secondary or tertiary structural changes. In contrast, single-domain proteins are almost completely packed, with no degrees of freedom for further compaction. The compact denatured structure induced by intramolecular aggregation may be ubiquitous in immunoglobulin proteins as non-canonical structures.

## Materials and Methods

Two monoclonal antibodies (mAb) commercially available as biotherapeutics were used; one is mAb-A, a humanised IgG1 monoclonal antibody (148 kDa; calculated pI^[66]^ = 8.5), and the other is mAb-B, a human IgG1 monoclonal antibody (147 kDa; calculated pI^[66]^ = 8.4). The molar mass includes the glycan mass. IgG1 was dissolved in a buffer solution (0.01 M sodium phosphate, 0.15 M NaCl and 0.005% (v/v) polyoxyethylene (20) sorbitan monolaurate, pH 7.4) to be ∼1–100 mg/mL (stock solution of IgG1). The IgG1 stock solution was diluted to the given concentrations (based on the experiment) with 0.01 M sodium phosphate buffer solution (pH 7.4) and dialysed against 0.01 M sodium phosphate buffer solutions at pH 3, 4, 6, and 7, or 0.1 M glycine-HCl buffer solutions at pH 1, 2, and 3 at 4 °C for 7 h using a Toru-kun TOR-14K microdialyzer (NIPPON Genetics Co., Ltd., Tokyo, Japan) with a membrane with a cut-off of 14 kDa. During dialysis and incubation, the protein aggregated at pH 1–2 (Figure S1). These samples were stored at 4 °C for ∼14 h until the batch SAXS measurement (Figure 1). Both the sodium phosphate and glycine-HCl buffer solutions at pH 3 produced comparable SAXS profiles (the latter is presented in Figure 1). For the SEC-SAXS measurement (Figure 2), the ∼10 or ∼20 mg/mL IgG1 stock solutions were dialysed against 0.1 M glycine-HCl buffer solution (pH 2.0) with a membrane cut-off of 12−14 kDa (Scienova GmbH, Jena, Germany) and stored at 4 °C for 1 h, 1 d, 3 d, or 7–8 d before measurement.

### SAXS

The SAXS experiments were performed with a BL-10C beam line at the Photon Factory of the High Energy Accelerator Research Organization (KEK) in Tsukuba, Japan.^[67]^ The X-ray wavelengths (*λ*) used were 0.10, 0.12, or 0.15 nm. The camera length was 1 or 2 m, and was calibrated using the scattering pattern of silver behenate.^[68]^ The scattering parameter *q* is defined as *q* = |***q***| = 4πsin*θ*/*λ*, where ***q*** is the scattering vector and 2*θ* is the scattering angle of the X-rays. X-ray intensities were recorded with a PILATUS3 2M detector (DECTRIS Ltd., Baden, Switzerland). For the batch SAXS measurement (Figure 1), the samples were measured at 25 ± 0.1 °C in a cell with quartz windows; 30 images were collected, where one image was recorded during a 2-s exposure time; the data from samples damaged by radiation were eliminated.

For the SEC-SAXS measurement (Figure 1), the IgG1 samples (∼7–11 mg/mL, 100–150 μL) were passed through a stainless-steel sample cell via an in-line size exclusion column (Superdex200™ (10/300) GL; GE Healthcare UK Ltd., Amersham, England; Lot. 10111292 and 10216439) operated by an HPLC system (Acquity UPLC H-class and Empower3 FR2; Waters Co., Milford, MA, USA; or Nexera-i; Shimadzu Co., Kyoto, Japan) installed with a beam line BL-10C^[67,69]^ and using flow rates of 0.2 mL min^−1^ for the initial 30 min and then 0.05 mL min^−1^ (Figure 2a, lower data) or 0.2 mL min^−1^ (Figure 2a, upper data). The temperature of the sample cell and the column was maintained at 25.0 °C. The exposure time for recording the image was 20 s. The sample solution was eluted into the sample cell with 0.02 mm-thick quartz glass windows and 1 mm-path length, where UV–vis absorption spectra (∼2 mm-path length by the diagonal incident light on the cell) were simultaneously recorded using a fibre-coupled UV–vis spectrometer, QEpro, with Opwave+ version 2.14–2.16 (Ocean Optics, Inc., Largo, FL, USA). A fitting analysis using theoretical scattering functions was conducted using the codes provided by the NIST Center for Neutron Research (NCNR).^[70]^ Unless otherwise indicated, all analyses were performed using IGOR Pro version 6.22A (WaveMetrics, Portland, OR, USA). SAXS measurements and the analyses of data are detailed in the Supporting Information and Table S4 therein.

### AUC

mAb-A samples at 0.7 mg/mL were used. Sedimentation velocity measurements were performed in a ProteomeLab XL-A (Beckman Coulter, Inc., Brea, CA, USA) with rotor speeds of 42 krpm at 25 ± 0.2 °C. An absorbance of 280 nm was used to monitor the protein concentration in a double-sector cell. The sedimentation velocity data were processed with the SEDFIT program using a sedimentation coefficient distribution analysis.^[71]^ The parameters (*s, f*/*f*_0_ and *f*/*f*_0,sp_) were determined according the equations in the Supporting Information.

### CD and IR

mAb-A samples at 1 mg/mL were measured. The procedures are detailed in the Supporting Information.

## Acknowledgements

This research is partially supported by the Japan Society for the Promotion of Science (grant numbers: JP19H03363 [to S.H.] and JP21K06503 [to H.I.]). The SAXS experiment in this work was performed under the approval of the Photon Factory Program Advisory Committee (Proposal No. 2015G061, No. 2017G068, and No. 2020G077). Computations were partially performed on the NIG supercomputer at ROIS National Institute of Genetics. The authors thank Dr. Tomoshi Kameda (AIST, Tokyo, Japan) and Dr. Seiki Yageta (AIST, Tsukuba, Japan) for stimulating discussions and comments.

## Author Contributions

A.I. and S.H. designed the research. H.I. and A.O. performed the experiments. H.I. analysed the data. H.I and S.H. wrote the main manuscript text. All authors contributed to data interpretation, critically reviewed the manuscript, and approved the final version of the manuscript.

## Data Availability

All study data are included in the article and/or *SI Appendix*.

