## Supplementary Information for "Getting Smaller by Denaturation: Acid-Induced Compaction of Antibodies"

Supporting information text (Results and Discussion)

1. pH-dependence of the small-angle X-ray scattering (SAXS) of mAb-A (Fig. S1)
2. Additional analyses of size exclusion chromatography small-angle X-ray scattering (SEC-SAXS) data for mAb-A at pH 2 (Fig. S2 and Table S1)
3. Analytical ultracentrifugation (AUC) for native and acid-denatured mAb-A (Fig. S3 and Table S2)
4. Circular dichroism (CD) and infrared (IR) spectroscopic analysis of native and acid-denatured mAb-A (Fig. S4 and Table S3)
5. Possible dimeric forms of acid-denatured mAb-A: a size exclusion chromatography small-angle X-ray scattering (SEC-SAXS) analysis (Fig. S5)
6. Theoretical aspect of anomaly of size-reduction upon acid-denaturation

Supporting information text (Materials and Methods)

SI References

**Supporting Information Text (Results and Discussion)**

**1. pH-dependence of the small-angle X-ray scattering (SAXS) of mAb-A**

The SAXS profiles of mAb-A at pH < 2.0 were distinct from those at pH 7.1–3.1, with a loss of the characteristic shoulders and peaks in the log–log plots and Kratky plots, respectively (Figure S1a and S1b, arrows). The significance of these scattering differences was also supported by the differences in the apparent radii of gyration ($R_g$) determined using Guinier plots (Figure S1c) and the apparent molar mass ($M$) calculated from the zero-angle scattering $I(0)$ (Figure S1d). The apparent $R_g$ and $M$ were close to the theoretical values at pH 7.1–3.1 and increased below pH 3.1, suggesting aggregation at pH 1.1 and 2.0. Indeed, the subsequent size-exclusion chromatography (SEC) analysis indicated protein aggregation at pH 2.0 (Figure 2a).

Accordingly, the distance distribution function ($P(r)$) at pH 1.1 indicated a larger maximum distance ($D_{max}$; Figure S1e), while $P(r)$ at pH 2.0 lacked two prominent peaks at pH 7.1–3.1 (Figure S1e, arrows). The peaks at the smaller $r$ (~40 Å) and larger $r$ (~80 Å) represented the intra- and inter-domain distance distributions of the electrons, respectively.[1,2] The mAb-A at pH 2.0 would therefore lose their native shape characteristics, such as the Fc (crystallizable fragment)-Fab (antigen-binding fragment) or Fab-Fab orientations. Negative $P(r)$ in the region of ~170 Å at pH 4.2 and 3.1 indicated an intermolecular interaction involved in the SAXS at a concentration of ~3.5 mg/mL.[3] The SAXS profiles at pH 7 (Figure S1f) and pH 3 (Figure S1g) depended on the protein concentration due to intermolecular interference.[4] The apparent $R_g$ (Figure S1h) and $I(0)/c$ (Figure S1i) indicated a larger concentration dependence at pH 3 than pH 7; thus, these results indicated that the effect of the interference led to a smaller apparent $R_g$ at pH 3.1 (Figure S1d). The second virial coefficients ($A_2$) at pH 7 and pH 3 were $(0.16 \pm 0.03) \times 10^{-4}$ and $(1.5 \pm 0.1) \times 10^{-4}$ cm$^3$ mol g$^{-2}$, respectively. Considering that the theoretical $A_2$ under hard sphere repulsion assumption (see

Materials and Methods in Supporting information) was $0.77 \times 10^{-4}$ $cm^3$ mol $g^{-2}$, the experimentally determined $A_2$ indicated an additional attractive interaction between mAb-A molecules at pH 7 and an additional repulsive interaction at pH 3, such as an electrostatic interaction. Aggregation at pH 2 also indicates that intermolecular attractive interactions should counteract the effect of the intermolecular repulsive interaction. Extrapolation of the apparent $R_g$ to zero concentration, which removes the interference effect, indicated similar $R_g$ values at pH 7 and pH 3. Moreover, all Kratky plots (Figure S1b) in the range of $q > 0.2$ Å$^{-1}$ represented the well-folded state of the mAb-A, which differs from the typical unfolded state model, such as a Gaussian coil.[3]

The statement, "the SAXS profiles were similar at pH 7.1–3.1" in the manuscript was judged by visual inspection of the common features of the scatterings at $q$ = ~0.04, ~0.08, and ~0.16 Å$^{-1}$, as highlighted by the Kratky plots (Figure 1b and Figure S1b). Nevertheless, the scatterings showed the variations in the peak positions and the intensities. We further conducted quantification of the similarity of the SAXS profiles. A pair-wise correlation map (CorMap) analysis[5] between the scattering at pH 7.1 and that at pH 6.2 indicated a $P$ value of 0.0454, thus, the hypothesis of similarity of the scattering data at pH 7.1 and that at pH 6.2 could not be rejected. However, CorMap analyses of all the remaining scattering data with the scattering at pH 7.1 presented $P < 10^{-6}$, thus, the hypothesis of similarity of the scattering data at pH 7.1 and that at an acidic pH condtion (pH 4.2, 3.1, 2.0, and 1.1) could be rejected, and the hypothesis of similarity of the scattering data at pH 7.1 and the theoretical scattering of IgG1 model (PDB: 1HZH) could also be rejected. Rejection of the hypothesis of similarity of the SAXS data could be due to conformational fluctuations of mAb-A rather than major conformational transition; e.g., CorMap analysis could discern scattering differences between open and closed forms.[5,6] We note that, prior to CorMap analysis, $I(q)$ was appropriately scaled to reduce the differences related to small uncertainities of the concentration determination. The data at $q$ = 0.017-0.512 was applied to CorMap analysis, which was conducted by *DATCMP* in the ATSAS 2.8.3 software.[7]

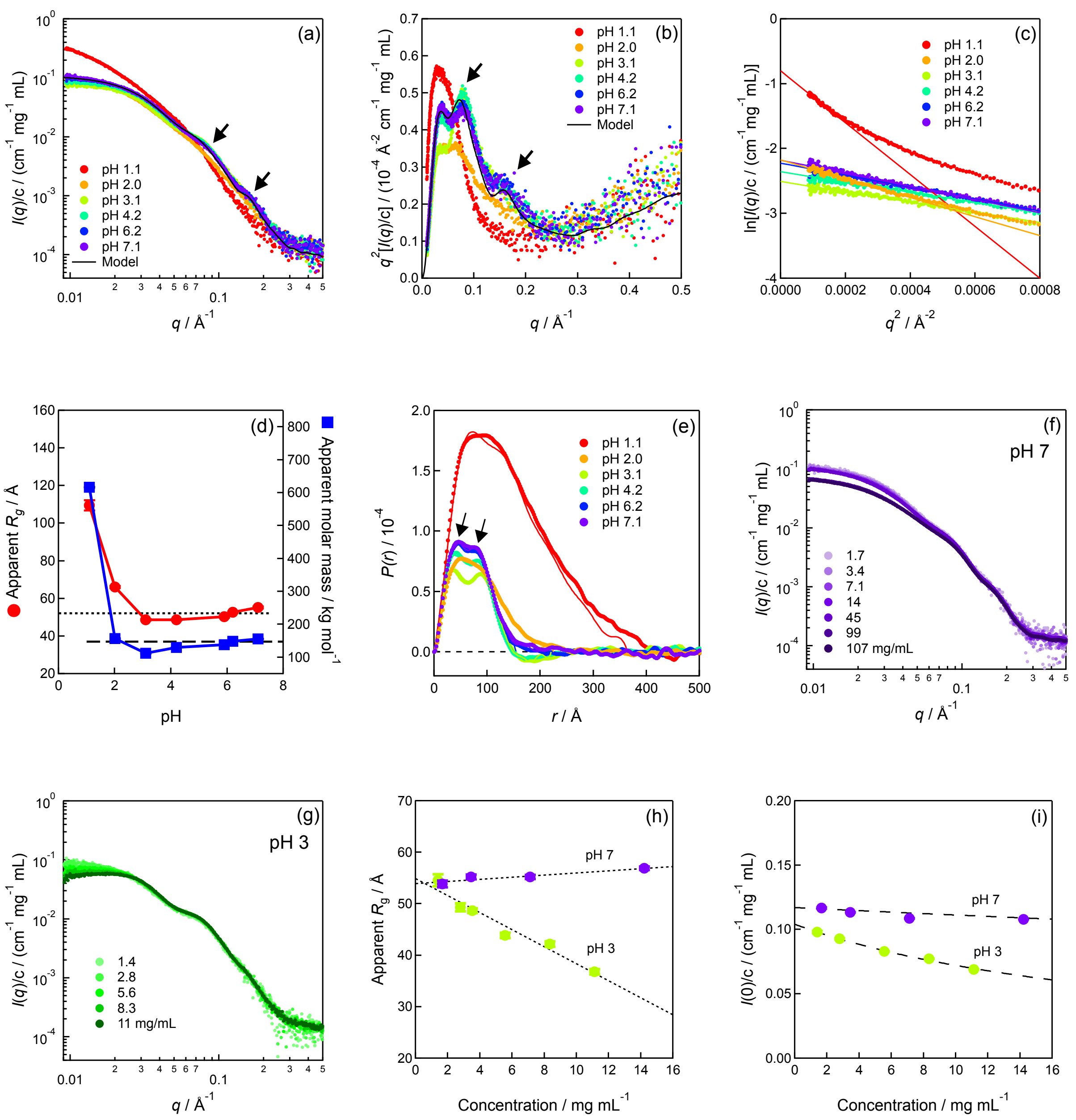


Figure S1. The mAb-A pH-dependence of the small-angle X-ray scattering (SAXS) at 25 °C. Concentration-normalized SAXS profiles are presented as (a) log–log, (b) Kratky, and (c) Guinier plots. The black lines in (a) and (b) show the theoretical SAXS profile of the IgG1 model (PDB:

1HZH). Solid lines in (c) indicate data fitted by Guinier approximation. (d) Apparent radius gyration ($R_g$) and apparent molar mass ($M$) determined by Guinier analysis; bars indicate standard error. We assumed $g = -1/3R_g^2$ at the finite concentration (equation S4) and $1 + 2A_2Mc_p = 1$ (equation S6), where $R_g$ and $M$ are apparent values (i.e., not the values extrapolated to zero concentration). (e) Distance distribution function, $P(r)$. The solid and dotted lines indicate indirect and direct Fourier transformations of the SAXS profiles, respectively. The protein concentration, $c$, in (a–e) was $3.5 \pm 0.1$ mg/mL and used for the normalization of $I(q)$. Concentration-dependence of the SAXS profiles at (f) pH 7 and (g) pH 3. Concentration-dependence of (h) $R_g$ and (i) $I(0)/c$ at pH 7 and pH 3 was calculated by Guinier approximation; error bars represent the uncertainty in the fitting analysis and were smaller than the marker size. Infinite dilution gives the exact $R_g$ and $I(0)/c$; otherwise, $R_g$ and $I(0)/c$ represent the effective values with the interference effect. Concentration-dependence of $R_g$ in (h) was approximated with a linear function, represented as dotted lines. Concentration-dependence of $I(0)/c$ (dashed lines) in (i) is described by equation (S6).

**2. Additional analyses of the size exclusion chromatography small-angle X-ray scattering (SEC-SAXS) data for mAb-A at pH 2.**

The frame-by-frame analyses of the SEC-SAXS data was used to inspect the stabilities of the $R_g$ (Figure S2a) and $I(0)/c$ (Figure S2b) values in each elution peak. For the elution peaks of the monomeric mAb-A ($R_g$ in Figure S2c and $I(0)/c$ in Figure S2d), Guinier plots of “Left”, “Top”, “Right”, and “Average” data fell on the same lines (Figure S2e and S2f), although the “Right” $R_g$ appeared to be bit smaller. The UV-chromatograms in Figure S2g indicate that mAb-A is more aggregation-prone at 0.2 M NaCl.

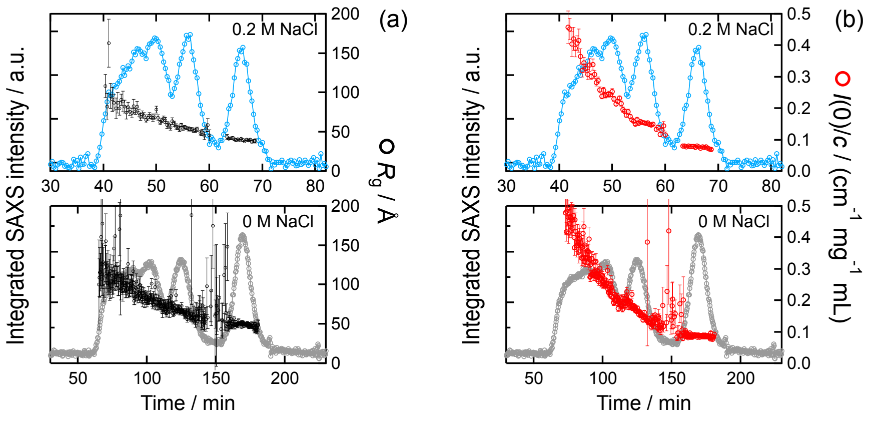


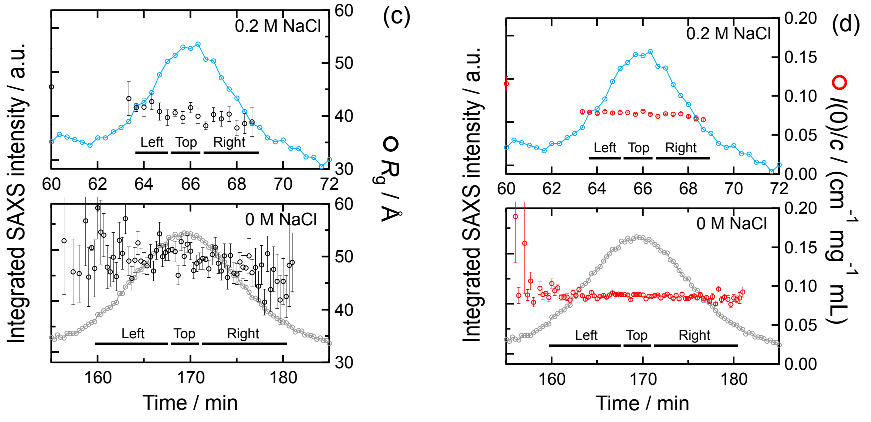

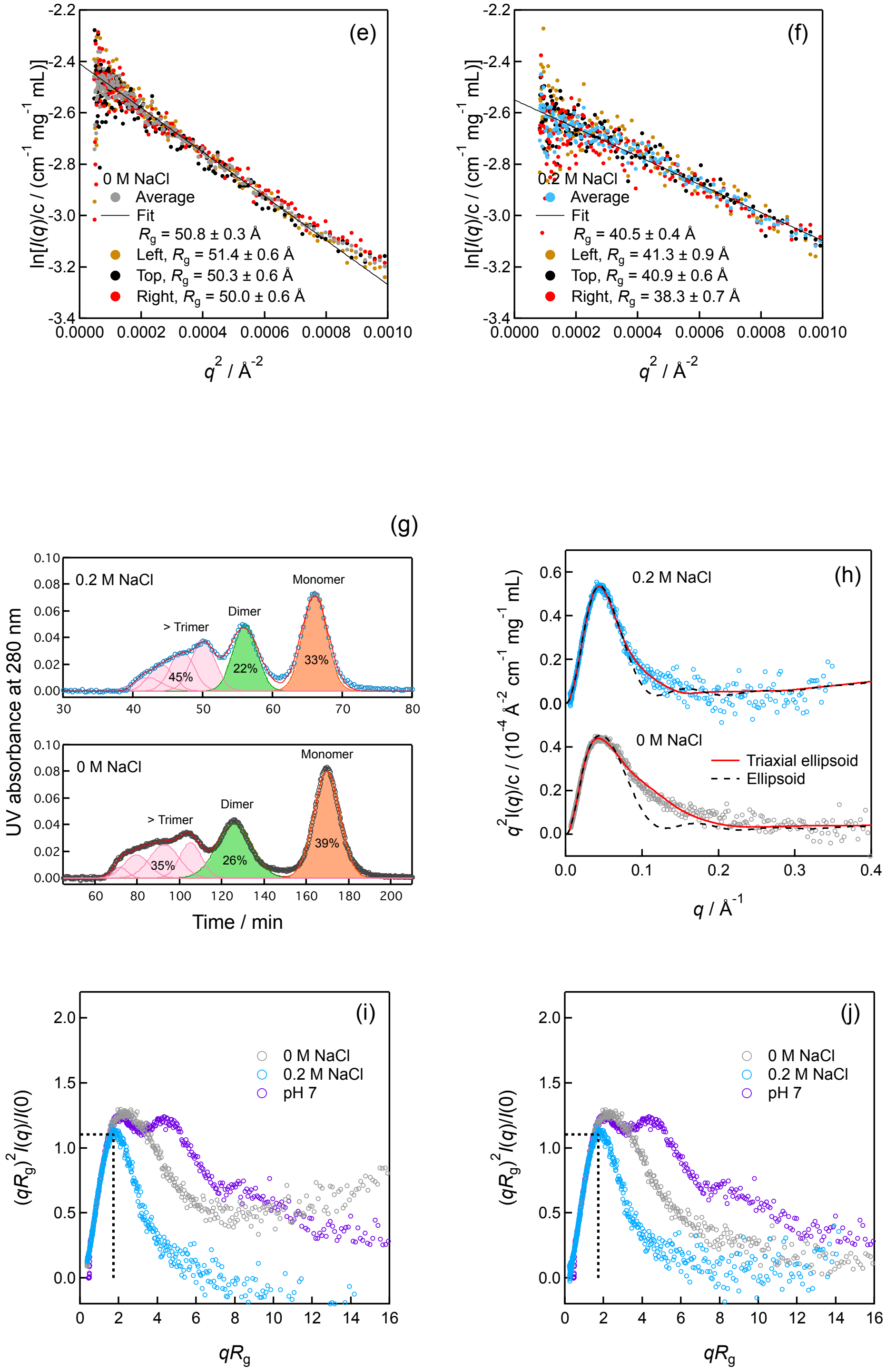


Figure S2. Additional analyses of the SEC-SAXS data for mAb-A at pH 2. Frame-by-frame analyses of (a) radii of gyration ($R_g$) and (b) $I/(0)/c$; $I(0)/c$ of the dimer was twice larger than that of the monomer. The elution peaks of the monomer in (a) and (b) are enlarged in (c) and (d), respectively. Guinier analyses of the data for the elution peaks of the monomer at (e) 0 M NaCl and (f) 0.2 M NaCl. “Left”, “Top”, “Right”, and “Average” represent that the SAXS data were averaged for the left side, the top, the right side, and the whole, respectively, of the elution peak in (c) and (d). The data of “Average” were used in the manuscript. (g) Peak area analyses of the chromatograms of the UV absorbance at 280 nm. The chromatograms were approximated as a sum of gaussian functions. The peak of the aggregates (> trimer) was assumed to be composed of four gaussian functions. (h–j) Kratky Plots of the monomeric mAb-A at pH 2. (i) and (j) present the dimensionless Kratky plots.[8,9] Porod correction (see Materials and Methods in Supporting information) was applied in (i), except for the data at pH 7. The black dotted lines indicate that the coordinate at $qR_g = 3^{1/2}$ and $(qR_g)^2I(q)/I(0) = 3\exp(-1) = 1.104$. (j) represents is the overlaid representation of Figure 2c.

Table S1. Molar mass and Porod volume analysis for mAb-A by the SAXS. The exact molar mass of mAb-A is 148 kg/mol.

| | | Molar mass / kg/mol | | Porod volume[c] / $10^3$ Å$^3$ |
|---|---|---|---|---|
| | | By $I(0)/c$[a] | By Bayesian inference[b] | |
| Sample | pH 7 | 121 | 157 (58%) | 234 |
| | pH 2, 0 M NaCl | 122 | 186 (88%) | 258 |
| | pH 2, 0.2 M NaCl | 108 | 186 (93%) | 239 |

[a]See Materials and Methods in the supporting information text. [b]Bayesian inference of the molar mass[10] (the probabilities are indicated in the parentheses) and [c]Porod volume calculations were conducted by the program *PRIMUS*[7] in the ATSAS 2.8.3 software.

### 3. Analytical ultracentrifugation (AUC) for native and acid-denatured mAb-A

Table S2. The sedimentation coefficient ($s$), friction factor ($f/f_0$), and corrected friction factor ($f/f_{0,sp}$) of the native and the acid-denatured mAb-A by the AUC

| Sample | $s$ / S[a] | $s$ / S (calculated) | $s_{20,w}$ / S | $f/f_0$ | $f/f_{0,sp}$[b] | $f/f_{0,sp}$ (calculated) | Duration of incubation |
|---|---|---|---|---|---|---|---|
| pH 7 (native) | 6.74 ± 0.06 | 6.35[c] | 6.66 ± 0.06 | 1.43 ± 0.01 | 1.20 ± 0.01 | 1.30[c] | — |
| pH 2, 0 M NaCl | 6.38 ± 0.07 | 6.86[d] | 6.06 ± 0.07 | 1.58 ± 0.02 | 1.32 ± 0.02 | 1.23[d] | 1 day |
| | 6.46 ± 0.11 | — | 6.13 ± 0.10 | 1.56 ± 0.03 | 1.31 ± 0.02 | — | 8 days |
| pH 2, 0.2 M NaCl | 7.02 ± 0.18 | 7.33[d] | 7.07 ± 0.19 | 1.35 ± 0.04 | 1.13 ± 0.03 | 1.09[d] | 1 day |
| | 7.00 ± 0.18 | 7.34[d] | 7.05 ± 0.18 | 1.36 ± 0.03 | 1.14 ± 0.03 | 1.08[d] | 8 days |

[a]Svedberg unit, S, $10^{-13}$ s. The errors of $s$ indicate the half-widths at half maxima of the fitted Gaussian distributions for the monomer components in $c(s)$ (Figure S3b), which are propagated in $s_{20,w}$, $f/f_0$, and $f/f_{0,sp}$ via equations S9–S12. [b]The values of $v$, $v_1$, and $\beta_1$ were used to calculate $f/f_{0,sp}$ (0.7425 $cm^3/g$,[11] 1.00296 $cm^3/g$,[12] and 0.51 g/g,[13] respectively). A $v$ for typical globular proteins was reported to decrease by ~0.4% upon complete unfolding.[14] Such small variations in $v$ from conformational changes can occur in the acid-denatured mAb-A and lead to slight changes in $f/f_{0,sp}$, but do not alter the present discussion and conclusion. [c]$s$ was simulated by the program

HYDROPRO[15,16] using the atomic coordinates for an IgG1 antibody (PDB: 1HZH). $f/f_{0,sp}$ was calculated from the *s* value according to equations (S9 and S10). These data are shown only as guides. $^{d}f/f_{0,sp}$ were calculated using the program ELLIPS[16,17] using the triaxial ellipsoid lengths, *a*, *b*, and *c*, as determined by the SAXS analysis in Figure 2. *s* was given according to equations S9 and S10.

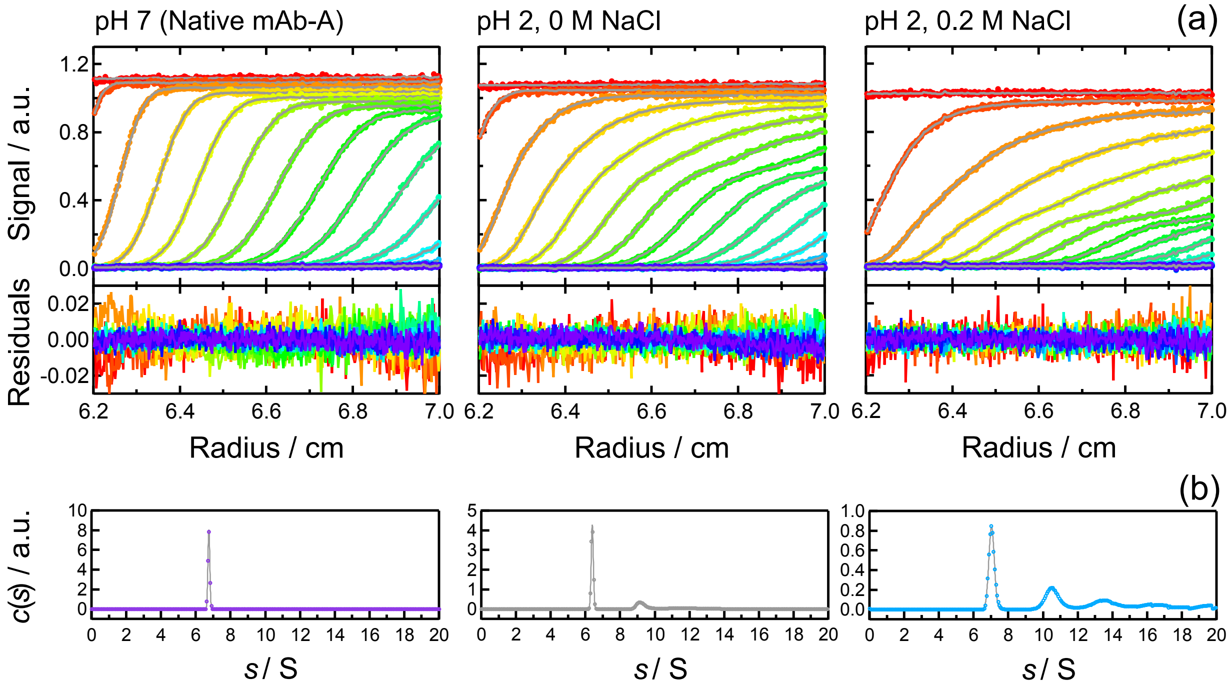


Figure S3. Analytical ultracentrifugation (AUC) for the native mAb-A and the acid-denatured mAb-A. (a) The sedimentation velocity data (closed circles), the fitted lines (grey solid lines), and their residuals for the proteins. (b) The sedimentation coefficient distributions, *c*(*s*), where the fitted lines were calculated by a Gaussian distribution function and indicate the monomer components. The acid-denatured mAb-A was incubated at pH 2 for 1 d prior to the measurements.

**4. Circular dichroism (CD) and infrared (IR) spectroscopic analysis of native and acid-denatured mAb-A**

The circular dichroism (CD) spectra captured secondary structural changes upon acid denaturation (Figure S4a), which inverted the sign of the ellipticities at ~200 nm from positive (pH 7, red) to negative (pH 2, grey), and a pronounced negative ellipticities at ~215 nm. Recent research has reported that twisting of β-sheets can generate a signal diversity at ~200 nm;[18,19] left-twisted and relaxed β-strands contribute to a positive signal, while right-twisted β-strands contribute to a negative signal. The analyses of the CD spectra using the BeStSel method[18,19] included these twisting effects (Table S3), indicating a decrease in the relaxed β-strands upon acid denaturation (Figure S4b).

The observed increase of the content of "others"[18,19] was consistent with the classical interpretation that describes an increase in the negative ellipticities at ~200 nm as a disordered structure,[20] the degree of which (5–9%) did not indicate a fully unfolded structure of the acid-denatured mAb-A. Rather, the secondary structures of mAb-A were marginally modulated upon acid denaturation. This β-rich character was supported by infrared (IR) spectroscopic analysis. The amide I bands stemming from the main chain amides were dominated by the antiparallel β-sheet signals, which typically emerged at ~1630–1640 $cm^{-1}$ and ~1680 $cm^{-1}$[21] for the native and acid-denatured mAb-A (Figure S4c). The peak frequency of the IR spectrum of the acid-denatured mAb-A (1632 $cm^{-1}$) was lower than that of the native IgG1 (1639 $cm^{-1}$), suggesting a minor alteration of the β-sheets, such as stronger hydrogen bonding within the β-sheets,[22] bifurcated hydrogen bonding of the main-chain amides with water,[23] or an untwisting of the right-handed twists of the antiparallel β-sheets.[24] The absorption at ~1728 $cm^{-1}$ for the acid-denatured mAb-A indicated a protonation of the side chains (-COOH).[21] The CD and IR spectra at pH 2 were measured without separation of the aggregates; prolonged incubation (12 d) at pH 2 caused a minor change in the CD spectra (difference between black and grey), most likely due to the mAb-A aggregation (Figure 2a).

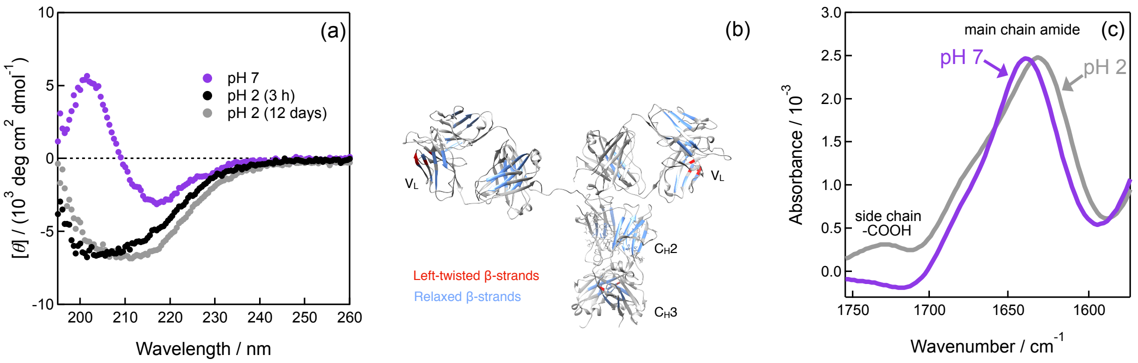


Figure S4. Spectroscopic analysis of the native mAb-A and the acid-denatured mAb-A. (a) Circular dichroism (CD) spectra of the native mAb-A (pH 7, red) and the acid-denatured mAb-A (pH 2). The acid-denatured mAb-A was incubated at pH 2 for 3 h (black) and 12 d (grey) prior to the measurements. The CD data are deposited in the Protein Circular Dichroism Data Bank (PCDDB), under the following accession codes: CD0006424000, CD0006424001 and CD0006424002. (b) Left-twisted (red) and relaxed (blue) β-strands mapped on the atomic coordinates of a crystal structure of IgG1 (PDB: 1HZH). The twisting of the β-strands was analysed using the program "HBSS_ext" in the SESCA software[25] and classified according to the definition of the BeStSel method.[18,19] (c) Infrared (IR) spectra of the native mAb-A (pH 7, red) and the acid-denatured mAb-A (pH 2, grey).

Table S3. Secondary structural contents (%) of native and acid-denatured mAb-A identified by CD spectra and BeStSel analysis[18,19]

| Secondary structure | pH 7 (native) | PDB: 1HZH[a] | pH 2 (3 h) | pH 2 (12 days) |
|---|---|---|---|---|
| Helix1 (regular) | 0.0 | 0.5 | 0.1 | 0.0 |
| Helix2 (distorted) | 0.0 | 2.7 | 2.8 | 0.9 |
| Anti1 (left-twisted) | 3.1 | 2.6 | 0.0 | 3.0 |
| Anti2 (relaxed) | 24.4 | 20.0 | 14.6 | 16.0 |
| Anti3 (right-twisted) | 20.9 | 18.9 | 19.2 | 17.9 |
| Parallel | 0.0 | 1.7 | 1.3 | 5.4 |
| Turn | 12.9 | 12.6 | 14.7 | 13.0 |
| Others | 38.7 | 40.9 | 47.3 | 43.7 |

[a]Secondary structural contents were calculated using the atomic coordinates of the IgG1 crystal structure (PDB: 1HZH). The positions of the left-twisted and the relaxed β-strands are mapped in Figure S4b.

### 5. Possible dimeric forms of acid-denatured mAb-A: A SEC-SAXS analysis

The ancillary benefit of the SEC-SAXS experiment was scattering from the separated aggregates. $R_g$ for the dimer of the acid-denatured mAb-A were 65.7 ± 1.0 Å at 0 M NaCl and 54.2 ± 0.8 Å at 0.2 M NaCl, which were comparable or smaller than the $R_g$ of native mAb-A dimers (e.g., a "compact" dimer = 64 ± 1 Å, and an "elongated" dimer = 78 ± 1 Å).[26] The experimental *P*(*r*) (Figure S5, pink circles) was comparable to the simulated *P*(*r*) of the two triaxial ellipsoids (black solid line), where the triaxial ellipsoids were oriented next to each other and slightly slipped. Face-to-face stacking of the triaxial ellipsoids (black dashed line) was not plausible due to the larger deviation from the experimental *P*(*r*).

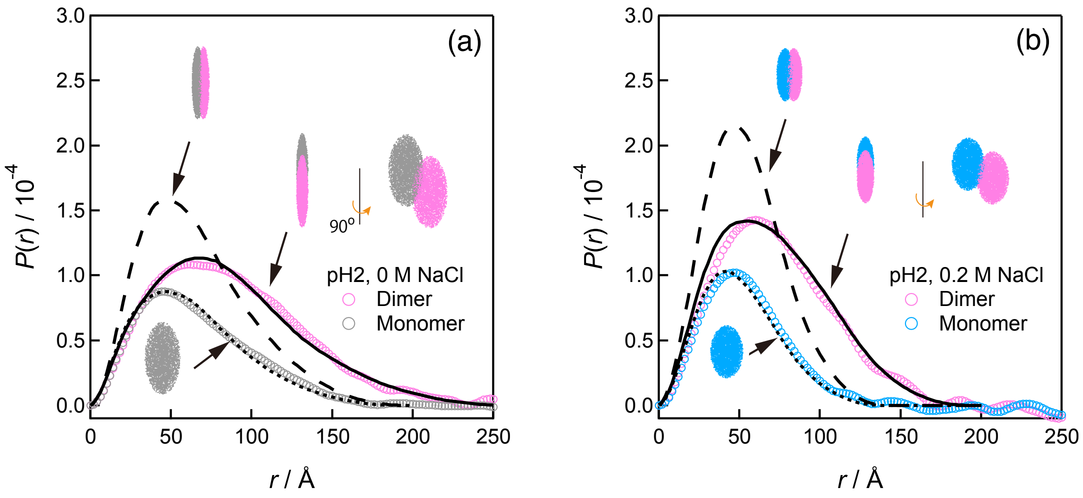


Figure S5. Possible dimeric forms of acid-denatured mAb-A. Pink and grey/cyan open circles indicate the *P*(*r*) of the dimer and the monomer, respectively, by direct Fourier transformation of the experimental SAXS profiles in Figure 2a at (a) pH 2 and 0 M NaCl (~125 min) and (b) pH 2 and 0.2 M NaCl (~55 min). The black dotted, dashed, and solid lines represent the simulated *P*(*r*) of the particles (arrows). The grey/cyan and pink dots illustrate the dimeric forms of the triaxial ellipsoids, which are the same size as the triaxial ellipsoid in Figure 2e. $P(r)$ values were approximated by the frequency function between the pairs of arbitrary points in the triaxial ellipsoid generated using the Monte Carlo method.[27]

### 6. Theoretical aspect of anomaly of size-reduction upon acid-denaturation: Linderstrøm-Lang smeared charge model.

When a spherical protein with an apparent radius ($r_{ap,j}$) and net charge ($Z_j$) is assumed to have evenly distributed charges on its surface, the electrostatic potential energy ($w(r_{ap,j})$) is given by[28,29]

$$w(r_{ap,j}) = e^2 l_D / [4\pi k_B T \varepsilon_{aq} r_{ap,j}(r_{ap,j} + l_D)] \qquad \text{(S1)}$$

where $e$, $l_D$, $k_B$, and $\varepsilon_{aq}$ are the elementary charge, Debye length, Boltzmann constant, and permittivity of water, respectively.[30] The subscript $j$ indicates the conformational state of the protein. For the transition from N (native) to D (denatured) states, the contribution of $w(r_{ap,j})$, which is denoted as $\Delta G_{elec}$, to the Gibbs energy for denaturation ($\Delta G_D$) is written as

$$\Delta G_{elec} = RT(w(r_{ap,D})Z_D^2 - w(r_{ap,N})Z_N^2) \qquad \text{(S2)}$$

Since $Z_N = Z_D$, the acid-denaturation ($\Delta G_{elec} < 0$) is accompanied by the size expansion, or $r_{ap,D} > r_{ap,N}$, which indicates that the size expansion decreases the Gibbs energy by denaturation.

**Supporting Information Text (Materials and Methods)**

**SAXS measurements** The circular 1D averaging of the images was performed using the *Nika* program.[31] The 1D scattering intensity data were averaged. The $q$-range available in the present study was ~0.01 < $q$ < 0.35–0.5 Å$^{-1}$. The scattering intensity was corrected based on the intensity of the incident light and the transmittance of the X-rays. The absolute scattering intensity of the protein ($I(q)$) (cm$^{-1}$) was determined by subtracting the corrected scattering intensity of the buffer solution ($I_B(q)$) from that of the protein sample solution ($I_S(q)$):

$$I(q) = [I_S(q) - (1 - c_p v)I_B(q)]/f_{cor} \quad \text{(S3)}$$

where $c_p$ is the protein concentration (g/cm$^3$), $v$ is the specific volume of the solute (cm$^3$/g), and $f_{cor}$ is the correction factor for converting the observed intensity in arbitrary units to the absolute intensity. A $v$ value of 0.7425 cm$^3$/g was used in this analysis, which is a good approximation for SAXS analysis of proteins.[11] The $f_{cor}$ values are dependent on the experimental setup and were determined by lysozyme scattering[11] for the SAXS data (Figure S1) and water scattering[32] for the SEC-SAXS data as standards. The lysozyme from hen egg white (Sigma-Aldrich, St. Louis, MO, USA) was dissolved in 0.01 M sodium phosphate buffer solution with 0.1 M NaCl (pH 7.4); the $c_p$ of the lysozyme was 0.0026 g/mL. The protein scattering in the low-$q$ region is approximated by:

$$\ln I(q) = gq^2 + \ln I(0) \quad \text{(S4)}$$

where $g$ is the slope, which gives a radius of gyration ($R_g$), or, $g = -1/3R_g^2$ at a diluted protein concentration according to Guinier law.[3] The $q$ regions appropriate for Guinier analysis were $q_{max}R_g < 1.3$, where $q_{max}$ is the largest $q$ for the regression, which was estimated using the program *AUTORG*[33] in the ATSAS 2.8.3 software.[7]

The distance distribution function, $P(r)$, was calculated by a direct Fourier transformation of $I(q)$ using the following equation:[3,34,35]

$$P(r) = \frac{1}{2\pi^2}\int_0^\infty I(q)qr\sin(qr)\exp(-Bq^2)\,\mathrm{d}q \qquad \text{(S5)}$$

where $r$ is the distance between the electrons in the particle and $B$ is the damping factor used to remove the termination effect of the Fourier transformation.[34,35] An indirect Fourier transformation was also performed using the GNOM program[36] in the ATSAS 2.8.3 software.[7] The maximum distance of the particle ($D_{max}$) where $P(r)$ approaches zero was estimated from $P(r)$. A theoretical SAXS profile of an IgG1 protein (PDB: 1HZH) was calculated using the atomic coordinates with the CRYSOL program.[37]

The absolute scattering intensity at $q = 0$, $I(0)$ is rationalised by the molar mass of the protein ($M$) using:

$$I(0) = kMc_p / (1 + 2A_2Mc_p) \qquad \text{(S6)}$$

where $k$ is a constant and $A_2$ is the second virial coefficient.[38,39] The $k$ value is equal to $[(1 - f_g)v(\rho_m - \rho_{solv}) + f_g v_g(\rho_g - \rho_{solv})]^2/N_A$, where $N_A$ is Avogadro's number, $(\rho_m - \rho_{solv})$ is the electron density difference between the protein and the solvent ($2.8 \times 10^{10}$ cm$^{-2}$),[11] and $(\rho_g - \rho_{solv})$ is the electron density difference between the carbohydrate and the solvent ($4.7 \times 10^{10}$ cm$^{-2}$).[40] $f_g$ is the weight fraction of the carbohydrate component (0.017 for mAb-A and 0.020 for mAb-B), and $v_g$ is the specific volume of carbohydrates (0.625 cm$^3$/g).[40] The value of $A_2$ for an ideal solution is 0, indicating there is no interaction between the particles. The value of $A_2$ for the repulsion between hard spheres with a radius $r_s$ (cm) is theoretically given by $2\pi N_A(2r_s)^3 / (3M^2)$.[39,41] The parameters approximated for mAb-A (148 kDa for $M$ and 55 Å for $r_s$) generated a hard sphere repulsion of $A_2 = 0.77 \times 10^{-4}$ cm$^3$ mol g$^{-2}$.

**SEC-SAXS measurements** Prior to injecting the sample, the solutions were filtered through a centrifugal filter unit with a pore size of 0.22 μm (Merck Millipore Ltd., Cork, Ireland) to remove possible large particle impurities.

SAXS-derived chromatograms were recorded as the SAXS intensities at all $q$ vs. retention time. The representative chromatograms are the integrated SAXS intensities over the $q$ range available vs. the retention time (Figure 2a). The scattering in the baselines of the chromatograms at $q$ were used as $I_\mathrm{B}(q)$; a linear increase of the baselines as derived from a capillary fouling[42] were corrected at all $q$. We determined the concentration-normalised $I(q)$, denoted as $I_\mathrm{c}(q)$ or $I(q)/c$, as follows: $I_\mathrm{c}(q)$ is given by $I_\mathrm{c}(q) = I_\mathrm{k}(q)/c_\mathrm{k}$, where $I_\mathrm{k}(q)$ is $I(q)$ at the concentration $c_\mathrm{k}$ for the frame number $k$. $I_\mathrm{k}(q)$ was determined using $c_\mathrm{k}$ and equation (S3). The average of $I_\mathrm{c}(q)$ can be calculated as $<I_\mathrm{k}(q)/c_\mathrm{k}>$, where the braces < > indicate an averaging over the frames. However, a simple application of this relation yields enhanced noise in $I_\mathrm{c}(q)$ because the smaller $c_\mathrm{k}$ amplifies the noisy $I_\mathrm{k}(q)$. Therefore, we used the summation of both sides of the relation $c_\mathrm{k}I_\mathrm{c}(q) = I_\mathrm{k}(q)$ such that $I_\mathrm{c}(q) = \Sigma I_\mathrm{k}(q)/\Sigma c_\mathrm{k}$, where $\Sigma I_\mathrm{k}(q)$ and $\Sigma c_\mathrm{k}$ were calculated independently to give the best quality. Due to the weak scatterings at the low protein concentration and potential flow-cell fouling issues[42], under- or over-subtraction of the background was seen (Figure S2i). This hinders the high-$q$ scattering-based analyses (e.g., structural heterogeneity, flexibility, and molecular weight estimation) but does not alter the low-$q$ scattering-based analyses (e.g., $R_\mathrm{g}$) or the main conclusion of this study. The under/oversubtraction was corrected by removing a small constant background from $I(q)$, assuming that the scattering at a higher $q$ obeys Porod's law: $I(q) \propto q^{-4}$.[3] This assumption holds when the scatterer is globular. The analytical ultracentrifugation indicates that the acid-denatured mAb-A is globular (see the Results in the manuscript).

Table S4. SAXS measurement parameters.

(*a*) Sample details

| | mAb-A | | | mAb-B |
|---|---|---|---|---|
| | pH 7 | pH 2, 0 M NaCl | pH 2, 0.2 M NaCl | pH 2, 0.2 M NaCl |
| Organism | *Mouse*/*human* (chimera) | | | *Human* |
| Source | Chinese hamster ovary cell-expressed (commercially available as biotherapeutics) | | | NS0 mouse myeloma cell-expressed (commercially available as biotherapeutics) |
| Extinction coefficient at 280 nm / ($cm^{-1}$ $M^{-1}$) | 204540 | | | 194300 |
| Partial specific volume / ($cm^3$ $g^{-1}$) | 0.7425[11] for the polypeptides<br>0.625[40] for the carbohydrates | | | |
| Average excess scattering density (contrast) / ($10^{10}$ $cm^{-2}$) | 2.8[11] for the polypeptides<br>4.7[40] for the carbohydrates | | | |
| Molecular weight from chemical composition / $10^3$ | 148 | | | 147 |
| SEC-SAXS | | | | |
| SEC column | | Superdex200™ (10/300) GL (Lot. 10111292) | Superdex200™ (10/300) GL (Lot. 10216439) | Superdex200™ Increase (10/300) GL (Lot. 10299331) |
| Loading concentration / mg $mL^{-1}$ | | 7 | 11 | 13 |
| Injection volume / μL | | 150 | 100 | 100 |
| Flow rate / mL $min^{-1}$ | | 0.2 (< 30 min)<br>0.05 (≥ 30 min) | 0.2 | 0.2 |
| Concentration (range) / mg $mL^{-1}$ | 1–107 | Avg. 0.29 (0.10–0.46) | Avg. 0.29 (0.12–0.41) | Avg. 0.68 (0.12–1.22) |
| Solvent details | 0.01 M sodium phosphate, pH 7 | 0.1 M glycine-HCl, pH 2 | 0.1 M glycine-HCl, 0.2 M NaCl, pH 2 | |
| Setup# | Setup#1,Setup#3 | Setup#2 | Setup#3 | Setup#4 |

Table S4 (continued).

(*b*) SAXS data collection parameters

| | Setup#1 (ID:20150515) | Setup#2 (ID:20160302) | Setup#3 (ID:20170305) | Setup#4 (ID:20211113) |
|---|---|---|---|---|
| Instrument | Beamline BL-10C with Dectris PILATUS 2M detector, the Photon Factory (PF) of the High Energy Acceleration Research Organization (KEK) | | | |
| Wavelength / Å | 1.488 | 1.000 | 1.200 | 1.200 |
| Beam size / $mm^2$ | 0.18×0.63 at focal point | | | |
| sample – detector distance / m | 1.047 | 2.006 | 2.028 | 2.081 |
| $q$-measurement range / $Å^{-1}$ | 0.0095–0.5780 | 0.0069–0.4259 | 0.0057–0.3486 | 0.0056–0.3396 |
| Absolute scaling method | Comparison with scattering from lysozyme | | Comparison with scattering from pure $H_2O$ | |
| Normalisation | To transmitted intensity by beam-stop counter | | | |
| Monitoring for radiation damage | Frame-by-frame comparison of data | | | |
| Exposure time | 2 s/frame (< 30 frames) | 2 s/frame (< 30 frames) for batch mode SAXS measurements<br>Continuous 20 s data-frame measurements of SEC elution for SEC-SAXS measurements | | |
| Sample configuration | cell with quartz windows, sample path length of 1 mm | | SEC–SAXS with flow cell, effective sample path length 1 mm | |
| Sample temperature / °C | 25 ± 0.1 | | | |

Table S4 (continued).

(*c*) Software employed for SAXS data reduction, analysis, and interpretation

| | |
|---|---|
| SAXS data reduction | *I(q) versus q* using *Nika*[31] |
| | solvent subtraction using *IGOR Pro* v.6.22A |
| Calculation of extinction coefficient from sequence | 5690 $cm^{-1}$ $M^{-1}$ for Trp and 1280 $cm^{-1}$ $M^{-1}$ for Tyr[43] |
| Basic analyses: Guinier, $P(r)$, and Porod volume | *PRIMUS* from *ATSAS* 2.8.3{Franke:2017hq} |
| Shape modelling | *IGOR* codes provided by the NIST Center for Neutron Research (NCNR)[44] |
| | *DENSS*[45] via web server (https://denss.ccr.buffalo.edu) |
| Atomic structure modelling | *CRYSOL*[37] in *ATSAS* 2.8.3 |
| 3D graphic model representations | *PyMOL* v.2.4.0a0 macOS |
| | *UCSF Chimera* v.1.13.1[46] |
| | *IGOR Pro* v.6.22A |

Table S4 (continued).

| (*d*) Structural parameters | | | | | |
|---|---|---|---|---|---|
| | mAb-A | | | | mAb-B |
| | pH 7 (1.7 mg/mL) | pH 7 (1.0 mg/mL) | pH 2, 0 M NaCl | pH 2, 0.2 M NaCl | pH 2, 0.2 M NaCl |
| | Setup#1 | Setup#3 | Setup#2 | Setup#3 | Setup#4 |
| Guinier Analysis | | | | | |
| $I(0)/c$ / $cm^{-1}$ | 0.1168 ± 0.0009 | 0.0880 ± 0.0007 | 0.0894 ± 0.0003 | 0.07834 ± 0.0004 | 0.1234 ± 0.0004 |
| $R_g$ / Å | 53.9 ± 0.7 | 50.3 ± 0.8 | 50.8 ± 0.3 | 40.5 ± 0.4 | 46.5 ± 0.3 |
| $q_{min}$ / $Å^{-1}$ | 0.0103 | 0.0081 | 0.0084 | 0.0109 | 0.0074 |
| $qR_g$ max | 1.3 | 1.3 | 1.3 | 1.3 | 1.3 |
| Coefficient of correlation, $R^2$ | 0.926 | 0.880 | 0.976 | 0.952 | 0.959 |
| Molecular weight from $I(0)$ / $10^3$ (ratio to expected value) | 160 ± 1 (1.08) | 121 ± 1 (0.82) | 123 ± 0.3 (0.83) | 108 ± 0.6 (0.73) | 169 ± 0.6 (1.15) |
| $P(r)$ analysis | | | | | |
| $I(0)/c$ | 0.1166 ± 0.0002 | 0.0875 ± 0.0003 | 0.08963 ± 0.0003 | 0.07812 ± 0.0004 | 0.1252 ± 0.0003 |
| $R_g$ / Å | 53.4 ± 0.1 | 50.6 ± 0.3 | 52.2 ± 0.2 | 40.8 ± 0.3 | 48.2 ± 0.1 |
| $D_{max}$ / Å | 156 | 149 | 176 | 142 | 166 |
| $q$ range / $Å^{-1}$ | 0.0102–0.1480 | 0.0081–0.1590 | 0.0083–0.2182 | 0.0109–0.1152 | 0.0074–0.1713 |
| Total estimate from *GNOM* | 0.83 | 0.89 | 0.78 | 0.85 | 0.87 |
| Molecular weight from $I(0)$ / $10^3$ (ratio to expected value) | 160 ± 0.3 (1.08) | 120 ± 1 (0.81) | 123 ± 0.3 (0.83) | 107 ± 0.6 (0.73) | 171 ± 0.4 (1.16) |
| Porod volume, $V_P$ / $10^3$ $Å^3$ (ratio /calculated molecular weight) | 236 (1.6) | 182 (1.2) | 259 (1.8) | 239 (1.6) | 237 (1.6) |

**AUC measurements** The ~5 mg/mL IgG1 (mAb-A) stock solution was dialyzed against 0.1 M sodium phosphate buffer solution (pH 7.2) and diluted to 0.7 mg/mL, after which it was referred to as the native IgG1 sample. To prepare acid-denatured IgG1 with/without 0.2 M NaCl, the ~10 mg/mL IgG1 stock solutions were dialyzed against 0.1 M glycine-HCl buffer solution (pH 2.0) with a membrane for cut-off of 12–14 kDa (Scienova GmbH, Jena, Germany) and stored at 4°C for 1 d or 8 d. One portion of the solution was diluted to 0.7 mg/mL IgG1 at pH 2 and 0 M NaCl, while another portion was dialyzed against 0.1 M glycine-HCl and 0.2 M NaCl buffer solution and diluted to 0.7 mg/mL IgG1 at pH 2 and 0.2 M NaCl. Prior to AUC, the solution was filtered through a centrifugal filter unit with a pore size of 0.22 μm.

The viscosity ($\eta$) of a 0.1 M sodium phosphate buffer solution (pH 7.2) at 25 °C was obtained (0.935 mPa s) using the SEDNTERP program.[47] The viscosities of 0.1 M glycine-HCl with/without 0.2 M NaCl buffer solution (pH 2.0) at 25 °C were estimated based on the assumption that the viscosity of a mixed solvent could be described by[48]

$$\eta/\eta_0 = 1 + \Sigma\Delta\eta_i \qquad \text{(S7)}$$

where $\eta_0$ is the viscosity of water at 25 ºC, 0.8905 mPa s.[47,49] $\Delta\eta_i = a_i c_i^{0.5} + b_i c_i$, where $a_i$ and $b_i$ are the coefficients in the Jones-Dole equation,[50] and $c_i$ is the concentration of the component $i$. $a_i$ and $b_i$ for glycine-HCl, HCl, and NaCl were determined by fitting the viscosity data at 25 °C (glycine-HCl) and at 20 °C (HCl and NaCl) reported in the literature[49,51] with the Jones-Dole equation in lower concentration regions. The temperature dependences of $a_i$ and $b_i$ were assumed to be negligible.

The sedimentation coefficient $s$ is represented by the re-established Svedberg equation:[52,53]

$$sRT/D = (1 - v_{sp}\rho_1)M_{sp} \qquad \text{(S8)}$$

where $R$, $T$, $D$, and $\rho_1$ are the gas constant, thermodynamic temperature, diffusion coefficient, and solvent density, respectively. The solvent densities were measured using a density meter DMA 5000 (Anton Paar, Graz, Austria). $M_{sp}$ is calculated by $M(1 + \beta_1)$, where $\beta_1$ is derived from hydration of the macromolecule; 0.30 g/g[53] and 0.51 g/g[13] for $\beta_1$ were reported. The subscript sp indicates the sedimenting particle. $v_{sp}$ is approximated by $(v + \beta_1 v_1)/(1 + \beta_1)$, where $v_1$ is the specific volume of the solvent.[52,53] $s$ is rationalised to the friction coefficient $f$ of the sedimenting particle as

$$f = (1 - v_{sp}\rho_1)(M_{sp}/N_A)/s \quad \text{(S9)}$$

The friction coefficient of a spherical particle, $f_{0,sp}$, with the equivalent volume of the sedimenting particle, is expressed as

$$f_{0,sp} = 6\pi\eta r_{0,sp} \quad \text{(S10)}$$

where $r_{0,sp}$ is the radius of the sphere, $[3v_{sp}M_{sp}/(4\pi N_A)]^{1/3}$, and $f/f_{0,sp}$ is the Perrin friction ratio function,[13] which provides information on how the shape of the sedimenting particle deviates from the sphere. The relationship between a classically employed friction factor, $f/f_0$, and $f/f_{0,sp}$ is expressed as[54]

$$f/f_{0,sp} = [(v + \beta_1 v_1)/v]^{-1/3} f/f_0 \quad \text{(S11)}$$

where $f_0$ is $6\pi\eta r_0$ and $r_0 = [3vM/(4\pi N_A)]^{1/3}$. $f/f_{0,sp}$ is treated as a corrected friction factor by incorporating the hydration effects missed in $f/f_0$. However, $f/f_0$ is often useful and superior to $f/f_{0,sp}$ when compared with previously reported values because $f/f_0$ is experimentally determined without the ambiguous parameter, $\beta_1$. $s$ at an infinite dilution of the protein ($s^0$) is corrected to $s_{20,w}$ as the sedimentation coefficient at 20 °C in water using:

$$s_{20,w} = s^0[(1 - v_{sp,20,w}\rho_{1,20,w})/(1 - v_{sp}\rho_1)](\eta/\eta_{20,w}) \quad \text{(S12)}$$

where $v_{sp,20,w}$ is $v_{sp}$ at 20 °C in water. $\rho_{1,20,w}$ is the density of water at 20 °C (0.998204 g/cm$^3$).[12] $\eta_{20,w}$ is the viscosity of water at 20 °C (1.002 mPa s).[47,49] Here, we assumed $v_{sp} = v_{sp,20,w}$ and $s = s^0$.

**Circular dichroism (CD) measurements** The native and the acid-denatured IgG1 (mAb-A) samples were prepared by dialyzing the ~10 mg/mL IgG1 stock solution against 0.01 M sodium phosphate buffer solution (pH 7) and 0.1 M glycine-HCl buffer solution (pH 2.0), respectively; and diluting the IgG1 solution to 1 mg/mL. The acid-denatured IgG1 was stored at 4 °C until the CD measurements. CD spectra were recorded using a J-805 CD spectrometer (JASCO Co. Ltd., Tokyo, Japan) with a temperature-controlled cell and a 0.02-cm path length (25 °C). The raw ellipticities were converted to mean residue ellipticities using the equation $[\theta] = \theta_{obs}/(10nc_Ml)$, where $\theta_{obs}$ is the observed ellipticity, $l$ is the optical path length (cm), $c_M$ is the concentration of the protein (molar), and $n$ is the number of residues.

**Infrared (IR) spectroscopic measurements** A 1 mg/mL IgG1 solution (pH 7.4) was used as the native IgG1 sample. The acid-denatured IgG1 samples were prepared by dialyzing the 11 mg/mL IgG1 stock solution against 0.1 M glycine-HCl buffer solution (pH 2.0) at 4 °C for 8 d and diluting it to 1 mg/mL. Infrared (IR) spectra were recorded using a Tensor 27 spectrometer (Bruker Optik GmbH, Ettlingen, Germany). The sample solutions were filled in a BioATR II attenuated total reflectance cell (Harrick, Pleasantville, NY, USA), which was connected to a thermostat (HAAKE K20, Thermo Electron Haake, Paramus, NJ, USA) set to 25 °C. A total of 128 interferograms were collected in a single beam mode with 4 cm$^{-1}$ resolution. The protein spectra were obtained by subtracting the corresponding buffer solution spectra.
